# Sleep promoting neurons remodel their response properties to calibrate sleep drive with environmental demands

**DOI:** 10.1101/2021.11.05.467459

**Authors:** Stephane Dissel, Markus K Klose, Bruno van Swinderen, Lijuan Cao, Paul J Shaw

## Abstract

Falling asleep at the wrong time can place an individual at risk of immediate physical harm. However, not sleeping degrades cognition and adaptive behavior. To understand how animals match sleep need with environmental demands, we used live-brain imaging to examine the physiological response properties of the *Drosophila* sleep homeostat (dFB) following interventions that modify sleep (sleep deprivation, starvation, time-restricted feeding, memory consolidation). We report that dFB neurons can distinguish between different types of waking and can change their physiological response-properties accordingly. That is, dFB neurons are not simply passive components of a hard-wired circuit. Rather, the dFB neurons themselves can determine their response to the activity from upstream circuits. Finally, we show that the dFB appears to contain a memory trace of prior exposure to metabolic challenges induced by starvation or time-restricted feeding. Together these data highlight that the sleep homeostat is plastic and suggests an underlying mechanism.

## Introduction

The importance of sleep is highlighted by the observation that it is evolutionarily conserved despite directly competing with all motivated waking behaviors [1, 2]. Not only does sleep compete with foraging, eating and mating [3–6] for example, high sleep drive may be maladaptive in many circumstances since falling asleep could place the individual in danger of immediate physical harm [7, 8]. On the other hand, sleep plays a critical role in learning and memory, supports adaptive behavior and facilitates creative insight. [9–12]. Together these observations suggest that it will not be possible to fully understand sleep’s function without knowing how sleep circuits calibrate sleep drive with motivational states.

In recent years, a great deal of progress has been made dissecting circuits that regulate motivated behavior in flies [13–17]. Typically, the properties of a circuit are examined as a function of one internal state. For example, sleep circuits are evaluated after sleep loss [18–20], feeding in response to starvation or high dietary sugar [21, 22], thirst following water deprivation [23], mating after social isolation [24] etc. However, since internal states can promote conflicting goal-directed behaviors, recent studies have begun to evaluate how circuits regulate competing activities such as feeding and sleep [5, 25–29], thirst and hunger [23], sweet and bitter taste [30], hunger and mating [31], and thirst vs hunger relevant memory [23], for example.

A common theme that has emerged from these studies has been that a specific deprivation-state differentially activates a subset of peptidergic neurons that then modulate classic neurotransmitter systems, frequently dopamine, to alter downstream circuits and thus motivated behavior [31–33]. For example, water deprivation preferentially activates a subset of peptidergic neurons which inhibit specific dopaminergic neurons to alter thirst-relevant memory [34]. In this model, the competition between state-specific peptidergic neurons will determine which motivated behavior will be expressed. An open question, however, is whether the neurons receiving peptidergic or dopaminergic input are constrained such that their responses are determined by upstream signals, or on the other hand, whether they can change their own response properties over time to influence a given outcome.

Sleep promoting *R23E10* neurons are well suited to investigate these relationships. *R23E10* neurons are modulated both by Dopaminergic neurons and Allatostatin-A (AstA)-expressing circadian-neurons [19, 23, 25–27, 35, 36]. AstA-expressing neurons promote sleep by releasing glutamate onto *R23E10* neurons [25]. Allatostatin-A is the *Drosophila* homologue of Galanin and has been implicated in both feeding and sleep [27, 37]. Sleep promoting *R23E10* neurons comprise a sleep switch and are an integral part of the sleep homeostat [19, 26]. The intrinsic properties of *R23E10* neurons have been evaluated in the context of sleep loss where it seems they monitor redox processes as an indicator of energy metabolism [19, 38, 39]. Thus, *R23E10* neurons may serve as command neurons of sorts that can integrate stimuli from competing internal states to gate information flow [40, 41]. Both the input and output connections to *R23E10* are well documented [25, 26, 35, 36]. Thus, the focus of this study is evaluate sleep promoting *R23E10* neurons following challenges that modulate motivational states.

In this study we identify independent sets of heterogeneous sleep-promoting neurons that change their responses to Dopamine and AstA following challenges that influence sleep. In addition, we identify a wake-promoting effect of AstA on *R23E10* neurons suggesting that the co-release of inhibitory AstA with excitatory glutamate may attenuate the overexcitement of *R23E10* neurons during high sleep drive and allow animals to maintain wakefulness in dangerous or life-threatening conditions. Finally, we find that both time-restricted feeding and acute starvation enhance subsequent waking by remodeling the expression of Dopamine receptors in sleep promoting neurons. Together, these data provide new insights into the interaction between internal states and sleep regulation.

## Results

### Neurons projecting to the dorsal Fan Shaped Body are modulated by Allatostatin

dFB projecting *R23E10* neurons are an important component of the sleep homeostat [19, 38, 39]. Sleep promoting dFB neurons are believed to be inhibited by wake-promoting dopaminergic neurons and activated when glutamate is released from sleep-promoting AstA expressing neurons [25, 35, 36]. Surprisingly, the role of AstA on *R23E10* neurons has not been investigated. Thus, we used behavioral genetics and live-brain imaging to characterize the effects of AstA and Dopamine on *R23E10* neurons. The expression pattern of *R23E10* neurons is shown in Figure 1a. Expressing the temperature-sensitive *Transient receptor potential cation channel* (*UAS-TrpA1*) in *R23E10* neurons and raising the temperature from 25°C to 31°C for 6 h increased sleep (Figures 1b,c). The parental controls do not show an increase in sleep during the exposure to 31°C (Figure S1). Since many neuromodulators act through second messenger signaling cascades, we expressed the cyclic adenosine monophosphate (cAMP) sensor, *UAS-Epac1-camps*, in *R23E10* neurons and used high throughput live-brain imaging to monitor neuronal responses to bath applied Dopamine (Figure 1d) [42, 43]. As seen in Figure 1e, DA increases cAMP levels in dFB neurons as previously reported [36]. It is important to note that while *R23E10>GFP* identifies ~14 neurons/hemisphere, *R23E10>UAS-Epac1-camps* reliably labeled ~8 neurons/hemisphere. Thus, we have confirmed that *R23E10* neurons are sleep promoting and respond to Dopamine.

**Figure 1:**
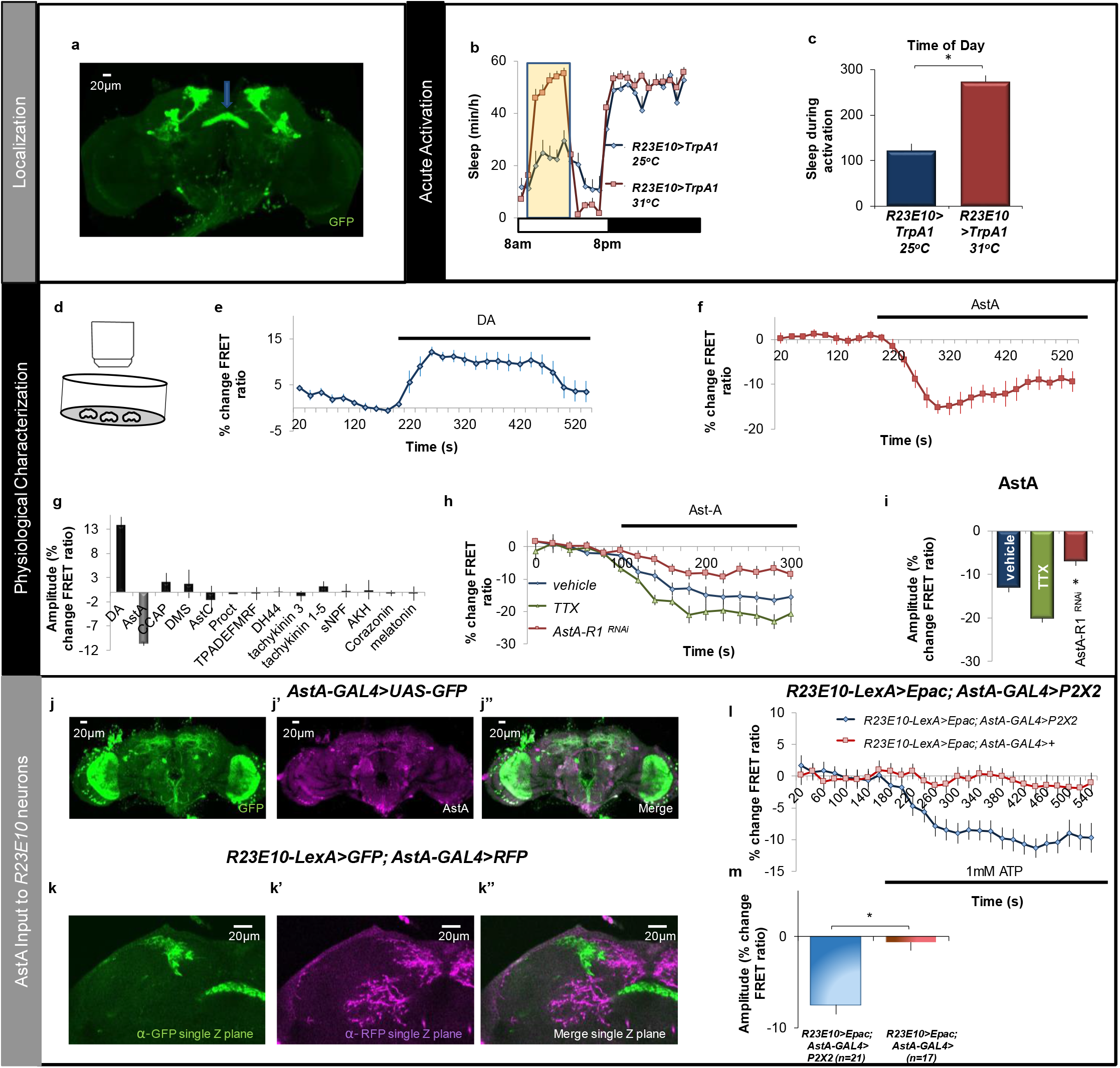
Sleep-promoting R23E10 neurons respond to AstA. **a)** Confirmation of *R23E10-GAL4>UAS-GFP* dFB projections (blue arrow). **b-c)** *R23E10-GAL4>UAS-dTrpA1* showed a significant increase in sleep at 31°C (red) compared to siblings maintained at 25°C (blue; n=16/group). **d)** Schematic of high-throughput Epac imaging. **e,f)** Normalized FRET ratio in *R23E10-GAL4>UAS-Epac1-camps* in response to DA (3 × 10^-5^ M) and AstA (1e^-6^M) (n=6-7 cells/condition). **g**) % change in FRET ratio in *R23E10-GAL4>UAS-Epac1-camps* following the application of neuroactive compounds (n = 6-21 cells/condition). **h)** TTX (1e-7M) does not prevent the response of *R23E10>UAS-Epac1-camps* to bath applied AstA (1e^-6^M) (n=4-6 cells/condition). Knocking down AstAR1 attenuates the response of R23E10 neurons to AstA (red trace; n=6). **i)** Quantification of H (p<0.05, modified Bonferoni test). **j)** Confocal stack of an *AstA-GAL4>UAS-GFP* fly brain stained with anti-GFP **j’)** Anti-AstA (magenta) antibody. **j”)** A merge image. **k)** Single optical section (0.5μm) of a *R23E10-LexA>LexAop-GFP, AstA-GAL4>UAS-RFP* fly brain stained with anti-GFP **k’)** Anti-RFP (magenta) **k’’)** A merge image. **l)** Application of 1 mM ATP induced a decrease in cAMP levels in *R2310* cells when the P2X2 receptor was expressed in *AstA-GAL4* expressing neurons (blue trace), but not in control lines lacking the P2X2 receptor (red trace). **m)** Quantification of l. Error bars represent standard error of the mean (SEM).

Given that *R23E10* neurons are downstream of AstA-expressing neurons [25], we evaluated the response properties of *R23E10* neurons to AstA and 14 other compounds that can influence motivational states [44]. As seen in Figure 1 f,g, AstA reduced cAMP levels in *R23E10* neurons. Previous studies indicate that AstA is an inhibitory peptide [45]. Individual traces for Figure 1g, are plotted in Figure S2a. Consistent with the cAMP responses, knocking down a battery of neuropeptide receptors in *R23E10* neurons using RNAi did not substantially modify sleep (Figure S2b). To determine whether the effects of AstA on *R23E10* neurons are direct, we incubated brains in Tetrodotoxin (TTX), which inhibits the firing of action potentials. TTX does not prevent the AstA mediated cAMP response of *R23E10* cells (Figure 1h, green trace). To further confirm a direct role of AstA we used RNA interference (RNAi) to knock down the *AstA-R1* receptor in *R23E10* neurons. As seen in Figure 1h (red trace), the cAMP response triggered by application of AstA is attenuated. Quantification of these effects are shown in Figure 1i. Importantly, *R23E10* neurons responded normally to DA when *AstA-R1* levels are knocked-down indicating that the reduced response to AstA is not the result of a non-functional neuron (Figure S2c).

To better understand the relationship between AstA and *R23E10* neurons we used immunohistochemistry to evaluate the overlap between AstA and *AstA-GAL4*. As previously reported, *AstA-GAL4* does not fully recapitulate the AstA expression pattern within the projections to the dFB (Figures 1j–j”) [37]. Nonetheless, AstA-GAL4 is expressed near the dendritic fields of *R23E10* suggesting a physical connection between the two group of neurons (Figures 1k-k”)[25]. To evaluate functional connectivity of the *AstA-GAL4, R23E10* circuit, we expressed the P2X2 activator [46] in *AstA-GAL4* neurons while measuring cAMP with *LexAop-Epac* in *R23E10-LexA* neurons. As seen in Figures 1l,m, perfusion of ATP which activates *P2X2*, leads to a reduction of cAMP levels in *R23E10* cells, mimicking the effect of AstA on *R23E10* neurons reported above (Figure 1f). No changes in cAMP signaling were observed in parental controls perfused with ATP but that do not express *P2X2* indicating the effects are specific to the activation of *AstA-GAL4* neurons (Figure 1l, red trace). Thus, we show that the cAMP response of *R23E10* neurons to AstA is similar to that seen when AstA expressing cells are activated.

The sleep promoting effects of AstA-expressing neurons have been shown to be due to the release of glutamate onto *R23E10* neurons [25]. Since AstA is an inhibitory neuropeptide [44, 45], we would predict the impact of AstA on *R23E10* neurons would be wake-promoting and that knocking down *AstARs* would thus increase sleep. To test this hypothesis, we evaluated sleep after knocking down *AstAR1* and *AstAR2* in *R23E10* neurons. As seen in Figures 2a,b and 2e,f, total sleep is increased when *AstA-R1* or *AstA-R2* are knocked down in *R23E10* neurons, suggesting that the dFB is under consistent allatostatinergic inhibitory tone. Importantly, sleep is also more consolidated during the day (Figures 2c,g); sleep consolidation is also increased at night (Figures S3a,b). Furthermore, the latency to fall asleep at night is reduced (Figures S3 c,d). The increase in sleep is not due to unhealthy or sick flies since waking activity is not reduced compared to parental controls (Figures 2d, h). To rule out off-target effects of the single RNAi line, we tested three additional, independent *AstA-R1* RNAi lines, and found that sleep is significantly increased in each line (Figure S3e). Given that increasing the activity of AstA neurons has been reported to increase sleep, but the *R23E10>AstAR^RNAi^* data indicate that AstA provides a wake signal, we asked whether our results might be due to unknown environmental factors in our laboratory. To evaluate this possibility, we utilized two lines (*AstA-GAL4* and *R65D05-LexA*) that express in AstA-expressing clock neurons and increase sleep [25]. As seen in Figure S3f, both *AstA-GAL4-dTrpA1* and *R65D05-LexA>LexAop-dTrpA1* lines substantially increased sleep consistent with previous reports [25]. Since AstA is an inhibitory neuropeptide, and the mammalian homologue of *AstA-R1* signals mainly through the Gi pathway [47], we hypothesized that the AstA receptors might be coupled to inhibitory G protein subunits. To test this hypothesis, we used RNAi to knockdown specific G protein subunits in sleep-promoting dFB neurons. As seen in Figure 2i, knocking down *Goα47A* had no effect on sleep while knocking down *Giα65A* using two different RNAi lines significantly increased sleep. Furthermore, independently knocking down β and γ1 subunits in sleep-promoting *R23E10* neurons also increased sleep (Figure 2i). Thus, these data suggest that AstA may be inhibiting dFB neurons via coupling with inhibitory G proteins.

**Figure 2:**
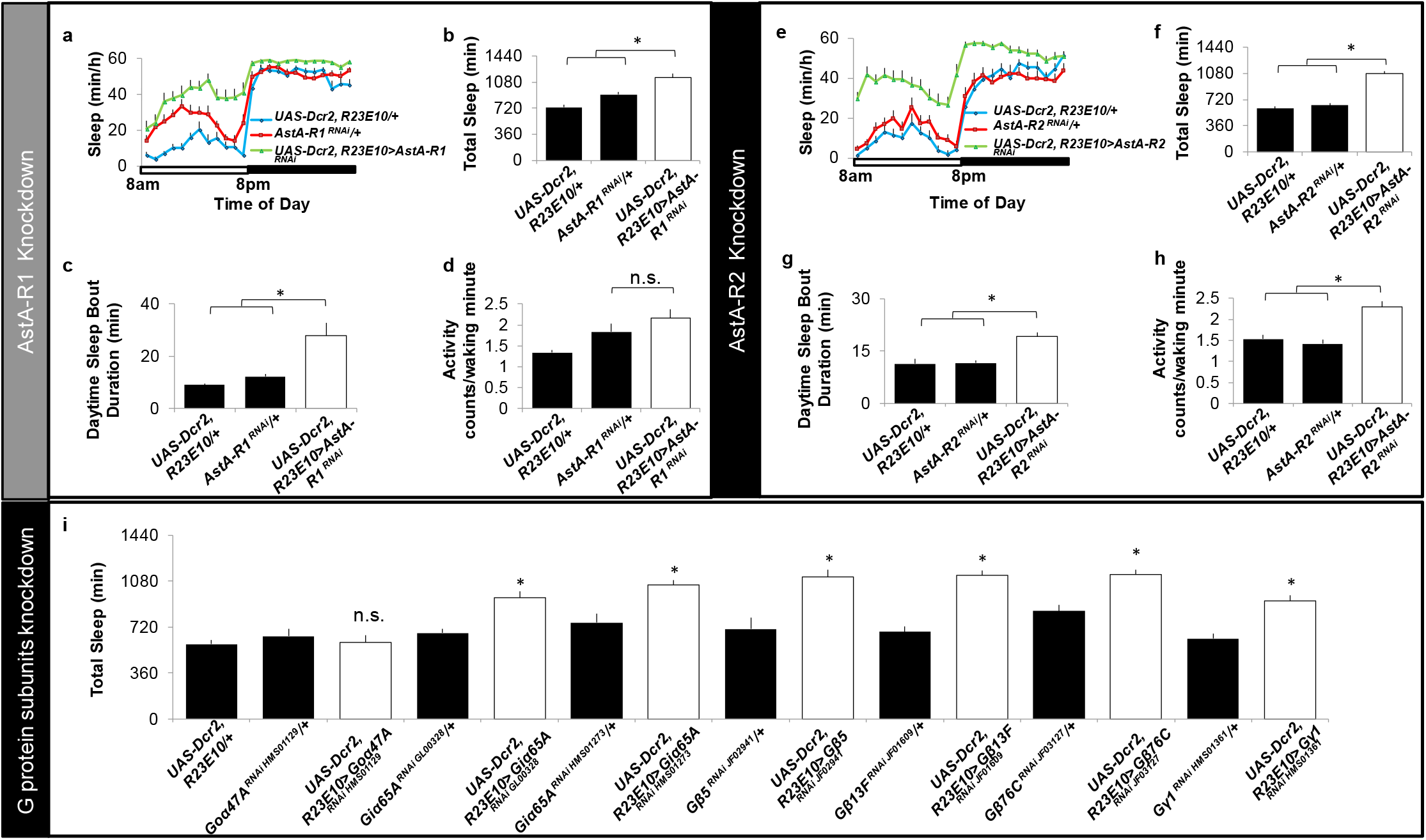
Knocking down *AstAR1* and *AstAR2* in *R23E10* neurons increases sleep. **a)** Sleep in minutes per hour **b)** Total sleep and **c)** daytime sleep-bout duration are increased when *AstA-R1* is knocked down in *R23E10* neurons compared with both *UAS-Dcr2, R23E10-GAL4/+* and *AstA-R1^RNAi^/+* parental controls (n=16/condition, *p<0.05 modified Bonferroni test). **d)** Waking activity is not different between *UAS-Dcr2, R23E10-GAL4/+> AstA-R1^RNAi^/+* experimental flies and both *UAS-Dcr2, R23E10-GAL4/+* and *AstA-R1^RNAi^/+* controls (n=16/condition, p>0.05, modified Bonferroni test). **e)** Sleep in minutes per hour **f)** Total sleep and **g)** daytime sleep bout duration are increased when *AstA-R2* is knocked down in *R23E10* neurons compared with both *UAS-Dcr2, R23E10-GAL4/+* and *AstA-R2^RNAi^/+* parental controls (n=16/condition, *p<0.05 modified Bonferroni test). **h)** Waking activity is significantly increased in *UAS-Dcr2, R23E10-GAL4/+> AstA-R2^RNAi^/+* experimental flies compared to both *UAS-Dcr2, R23E10-GAL4/+* and *AstA-R2^RNAi^/+* controls (n=16/condition, p<0.05, modified Bonferroni test). **i)** Total sleep is not changed when Goα47A levels are reduced in *R23E10* neurons, but independently knocking down *Giα65A, Gβ5, Gβ13F, Gβ76C* and *Gγ1* in *R23E10* cells increase total sleep compared with parental controls (n=16/condition; *p<0.05). Error bars represent SEM.

### Sleep promoting dFB are diverse

Dorsal Fan Shaped Body neurons (dFB) have been hypothesized to gate different aspects of sleep such as locomotion, sensory thresholds etc. [26, 48]. This hypothesis suggests that the dFB could be comprised of independent sets of sleep promoting neurons that each respond to distinct environmental challenges. If this were to be the case, then we should be able to identify novel dFB projecting GAL4 lines. To expedite the identification of such lines, we asked Dr. Gerry Rubin and Dr. Arnim Jenett, for assistance [49]. They hand-selected 12 *GAL4* lines that matched the expression pattern of the original sleep promoting GAL4 lines (*104y* and *C5*) [10]. We activated these neurons by expressing *UAS-TrpA1* as described above. Surprisingly, only one driver, *R55B01* increased sleep similarly to *R23E10* when compared to siblings maintained at 25°C (Figures 3 a,b and Figure S4). The parental controls do not show an increase in sleep during the 6h exposure to 31°C (Figure S1). Similar results were obtained when expressing the sodium bacterial channel, *UAS-NaChBac* (data not shown). As seen in Figure 3c, *R55B01* project to the dFB neuropil (blue arrow) and have cell bodies located in the same anatomical region in the brain as *R23E10* neurons. To determine potential overlap between *R23E10* and *R55B01*, we conducted co-labeling experiments to express RFP with GAL4/UAS and GFP with LexA/LexAop in the same fly. As seen in Figures S5c-c”,d *R23E10-LexA* is similar, but not identical, to the expression pattern of *R23E10-GAL4*. Importantly, *R23E10-LexA* and *R55B01*-GAL4 only share ~4 neurons in common (Figures S5 a-a”,b). As with *R23E10* neurons, *R55B01> UAS-Epac1-camps* labels fewer dFB projecting neurons (~8 neurons) compared to *R55B01>GFP*. Similar to *R23E10* neurons, *R55B01>UAS-Epac-camps* show robust responses to AstA (Figure 3d). Moreover, AstA positive staining is co-localized with the GFP positive processes of *R55BO1* neurons (Figures S5e-e””). In contrast to *R23E10* neurons, the response of *R55B01* neurons to dopamine is either reduced or absent (Figure 3e). A box plot comparing the range of responses of *R23E10* and *R55B01* neurons to dopamine is shown in Figure 3f. Given the robust responses of *R55B01* neurons to AstA, we evaluated sleep after knocking down *AstAR1* and *AstAR2*. As seen in Figures 3g,h sleep is increased upon knocking down either *AstAR1* or *AstAR2* in *R55B01* neurons. Quantification of sleep architecture in *R55B01>AstAR1^RNAi^* and *R55B01>AstAR2^RNAi^* can be found in Figure S6. Thus, we have identified an additional sleep promoting driver that projects to the dFB, has little overlap with *R23E10* neurons and is less responsive to dopamine.

**Figure 3:**
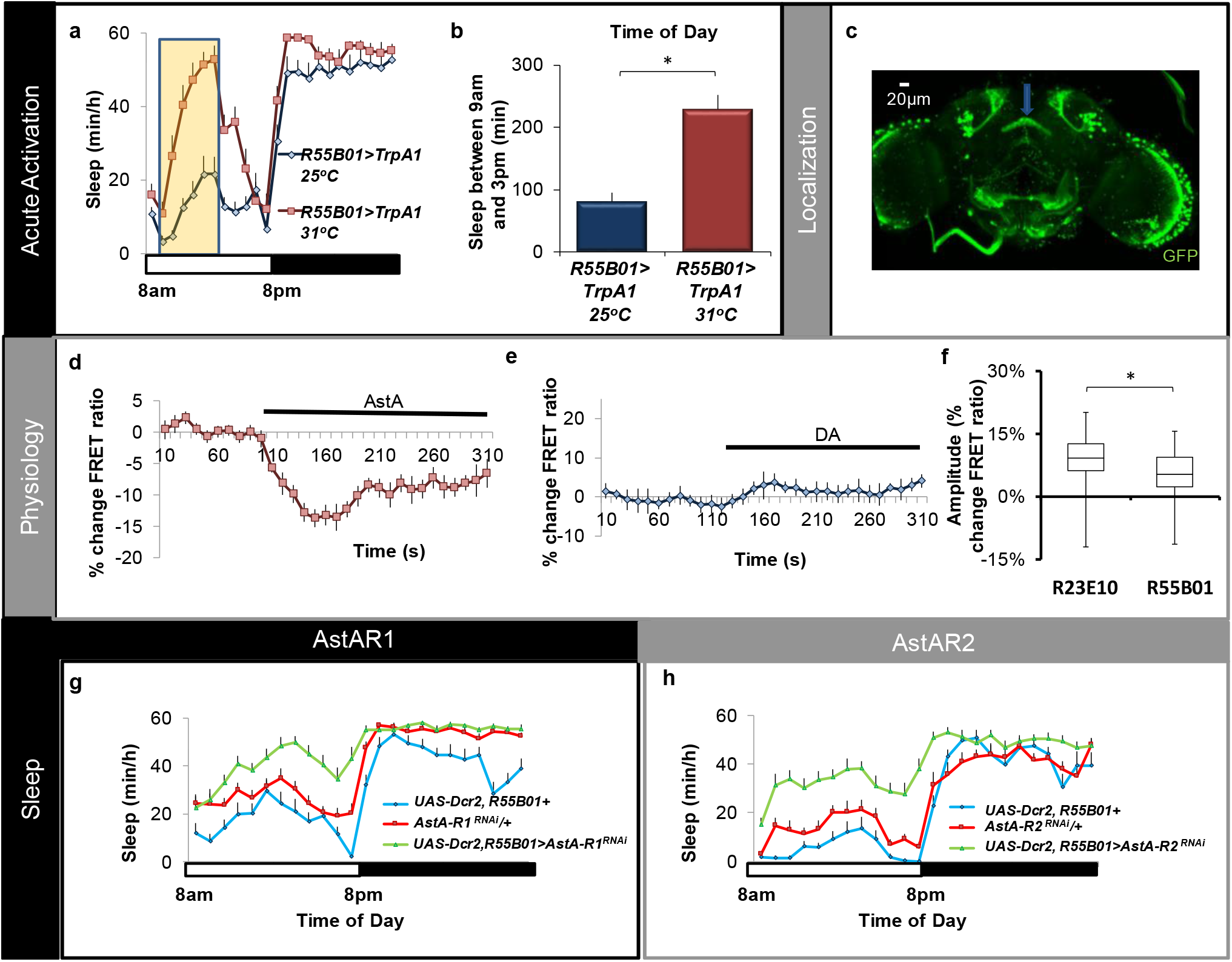
R55B01 neurons respond to Allatostatin-A. **a-b)** *R55B01-GAL4>UAS-dTrpA1* flies showed a significant increase in sleep at 31°C (red) compared to siblings maintained at 25°C (blue; n=16/group). **c)** Expression pattern of *R55B01-GAL4>UAS-GFP* (blue arrow indicates the location of dFB). **d)** Normalized FRET ratio in *R55B01>UAS-Epac1-camps* in response to 1e^-6^M AstA (n = 23 cells). **e)** Normalized FRET ratio in *R55B01>UAS-Epac1-camps* to 3e^-5^M DA (n = 10). **f)** Box plot comparing the response properties of *R23E10>Epac* and *R55B01>Epac* neurons to 3e^-5^M DA. The bottom and top of each box represents the first and third quartile, and the horizontal line dividing the box is the median. The whiskers represent the minimum and maximum individual cell responses.n=68 cells for R23E10 and n=115 cells for R55B01. **g,h)** Knocking down *AstAR1* or *AstAR2* in R55B01 neurons increases sleep; Sleep in minutes per hour in *R55B01-GAL4>AstAR1^RNAi^* and *R55B01-GAL4>AstAR2^RNAi^* (n=16/condition). Error bars represent SEM.

### R23E10 neurons differentially integrate sleep-relevant stimuli

Together with the literature, these data identify two major wake-promoting signals that impact *R23E10* neurons; AstA and Dopamine [35, 36]. Although both signals are inhibitory, they derive from neuronal circuits with opposite functions; AstA is released from sleep promoting neurons while Dopamine is released from wake-promoting neurons [25, 35, 36]. Given these divergent roles, we hypothesize that AstA and Dopamine will operate largely independently. To test this hypothesis, we used the live-brain imaging approach described above (Figure 1d) to evaluate the response of *R23E10* neurons to either Dopamine or Allatostatin during conditions that alter sleep drive. Since *R55B01* neurons respond less to Dopamine, we focused solely on *R23E10* neurons; *R55B01* neurons will be studied later. We first evaluated the response properties or *R23E10* neurons in unperturbed flies under conditions characterized by large changes in sleep time (e.g. ontogeny, gender, individual differences). We then evaluated the response properties of *R23E10* neurons after experimental interventions that modulate sleep drive (e.g. sleep loss, starvation, training that induces long-term memory) [3, 28, 50–53] (Figure 4). Quantification of traces in Figure 4 can be found in Figure S7).

**Figure 4:**
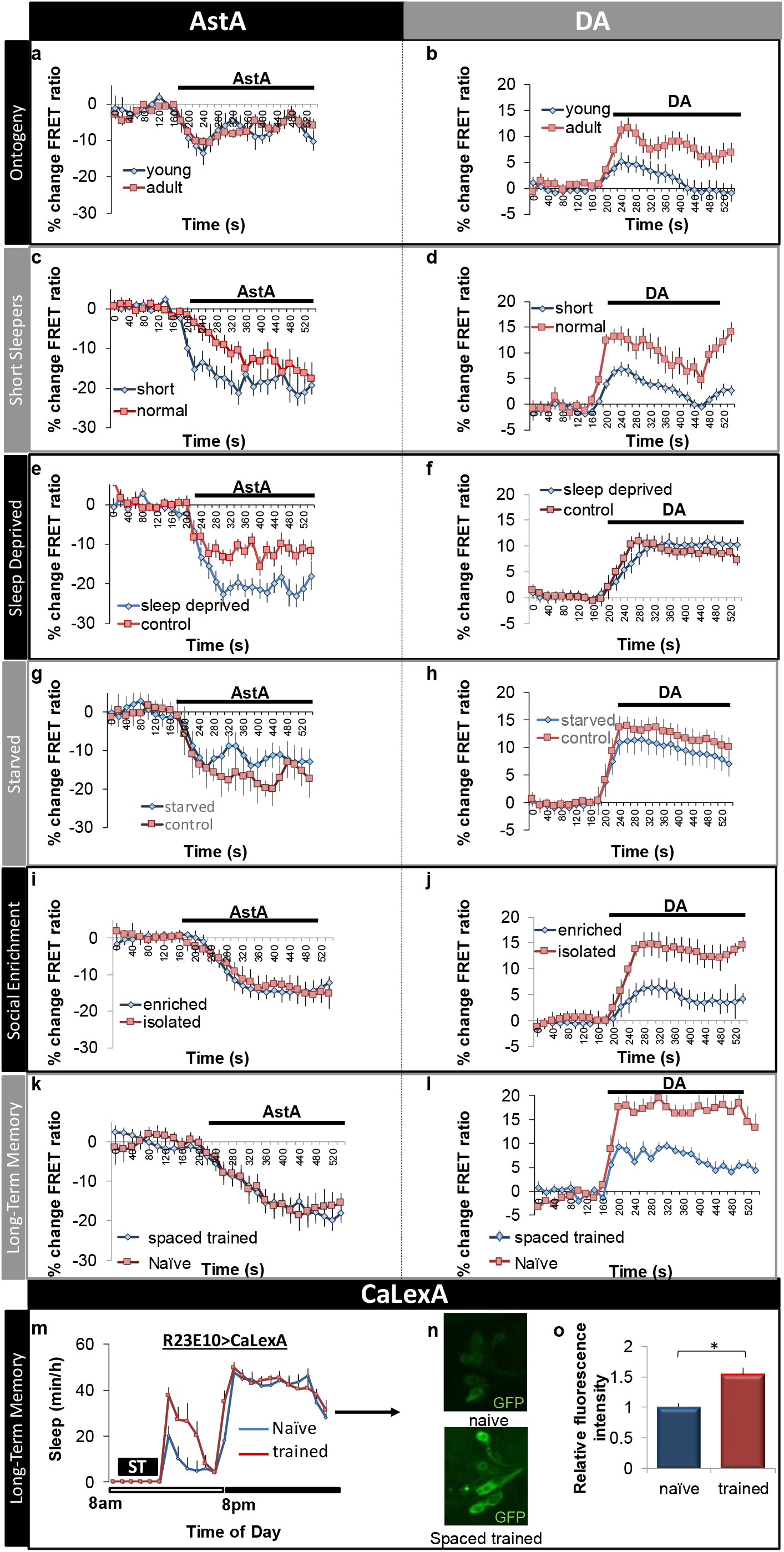
Sleep-promoting neurons integrate a variety of relevant stimuli. **a,b)** Response of *R23E10* neurons in young (0-1 day old) and adult (6-8 day old) flies in response to Allatostatin-A (10^-6^M) (n = 13, 22 cells) or Dopamine (3e^-5^M) (n = 15,14 cells). Quantification of traces in Figure 4 can be found in Figure S7. **c,d)** Response of *R23E10* neurons in spontaneously short sleeping flies (<400min/day) and normal sleeping siblings (600-900min/day) (AstA, n = 18,20 cells) and (Dopamine, n = 12,19 cells). **e,f)** Response of *R23E10* neurons following 12 h Sleep deprivation compared to untreated siblings (AstA, n = 17,17 cells) and (Dopamine, n = 29,26 cells). **g,h)** Response of *R23E10* neurons following 18 h starvation compared to untreated siblings; (AstA, n = 20,14 cells) and (Dopamine, n = 23,16 cells). **i,j)** Response of *R23E10* neurons following social enrichment compared to isolated siblings;(AstA, n = 9,9 cells) and (Dopamine, n = 12,8 cells). **k,l)** Response of *R23E10* neurons following spaced training compared to naïve controls (AstA, n = 20,10 cells) and (Dopamine, n = 25,10 cells).**m)** Sleep is increased in *R23E10-GAL4>CaLexA* flies following a spaced courtship training (n=10 per group, p<0.05). **n)** Representative confocal stack of the brains of a naïve (top) and spaced trained (bottom) *R23E10-GAL4>CaLexA* flies stained with anti-GFP antibody. Flies were dissected on the day after courtship conditioning. **o)** Quantification of GFP staining intensity in R23E10 neurons in naïve and spaced trained *R23E10-GAL4>CaLexA* flies (n=107 neurons for naïve and 100 neurons for trained).

As with humans, sleep is highest in young flies and then stabilizes in early adulthood. Interestingly, *R23E10* neurons are less responsive to Dopamine in 0-1day old flies compared to 6-8 day old mature adults; no changes were observed in response to AstA (Figures 4a,b). The response properties of *R23E10* neurons to Dopamine were similar in 6-8 day old flies and 30-38 day old flies (Data not shown). In contrast to age, we did not observe any changes in the response properties of *R23E10* neurons in male and female flies despite large sexual dimorphisms in sleep behavior [52, 54] (Data not shown). We have previously shown that individual differences in sleep time can be exploited to evaluate sleep regulation and function [55]. Importantly, our previously published data indicate that spontaneously short-sleeping flies appear to experience both high wake-drive and high sleep-drive simultaneously [56]. With that in mind, we evaluated the response properties of *R23E10* neurons in spontaneously short-sleeping *R23E10>UAS-Epac-camps1*(*i.e*. total sleep time less than 400min) compared to normal sleeping siblings (800-960 min sleep). As seen in Figure 4c, *R23E10* neurons appear to be under stronger inhibitory tone from AstA in spontaneous short-sleepers compared to normal sleeping siblings. Although one would predict a stronger response of *R23E10* neurons to the wake-promoting effects of Dopamine, short sleeping flies were less responsive to Dopamine (Figure 4d). Future studies will be needed to determine whether incongruent responses of *R23E10* neurons to AstA and Dopamine reveal underlying deficits in sleep regulation.

We next asked whether distinct experimental interventions that modulate sleep drive would alter the physiological response properties of *R23E10* neurons in similar or dissimilar ways. Both sleep deprivation and extended periods of starvation are followed by a compensatory increase in sleep [3, 28, 57]. To evaluate how sleep disruption would influence *R23E10* neurons, flies were sleep deprived for 12 h or starved for 18 h and compared to untreated siblings. As seen in Figure 4e, cAMP responses to AstA are increased following sleep deprivation but were unchanged in response to Dopamine (Figure 4f). Surprisingly, the response of *R23E10* neurons to either AstA or Dopamine were not changed while being starved (Figures 4g,h and see below). In contrast to sleep deprivation and starvation, social enrichment induces plasticity in specific neural circuits to increase sleep without exposing flies to sleep loss [51, 58]. Changes in sleep following social enrichment have been mapped to pigment dispersing factor (PDF) expressing clock neurons [58]. Interestingly, AstA signaling is modulated by PDF [27]. Thus, to evaluate the impact of social rearing we housed flies in a socially enriched environment (50 flies/vial) for 5 days and evaluated cAMP responses compared to isolated siblings. As seen in Figure 4i, *R23E10* neurons of socially enriched or isolated flies show similar responses to AstA. However, DA responses of *R23E10* cells are strongly reduced by social enrichment (Figure 4j). These data demonstrate that disparate behavioral manipulations that increase sleep drive in different ways induce independent physiological responses of *R23E10* neurons to AstA and DA.

Sleep dependent memory consolidation has been linked to both the ventral and dorsal Fan Shaped Body [10, 59]. Thus, we asked whether *R23E10* neurons would modify their physiological responses to a training protocol that induces LTM [9]. As seen in Figure 4k, responses of *R23E10* neurons to AstA are not different following courtship conditioning. However, training resulted in a dramatic reduction in the response of *R23E10* neurons to DA (Figure 4l). Interestingly, the effect of training on DA responses could still be observed 24h after the end of the training only returning to baseline 48h later suggesting long-term plastic changes in *R23E10* neurons (Figure S7m). To determine whether the changes in cAMP were due to non-specific effects of courtship or to memory consolidation, we evaluated cAMP levels following a massed training protocol consisting of a single 3h session that does not result in LTM formation [10]. As seen in Figure S7m, massed training did not alter the responses of R23E10 cells to Dopamine. Furthermore, no changes in responses to Dopamine were found when training was followed by 4 h of sleep deprivation (data not shown). To further evaluate the role of the dFB in courtship memory, we used *CaLexA* (*calcium-dependent nuclear import of LexA*) to see if *R23E10* neurons might show sustained activity following a spaced training protocol [60]. *R23E10* flies expressing CaLexA were exposed to a training protocol consisting of 3×1h individual pairings of a naïve male with a non-receptive female target separated by a rest period of 1h. As seen in Figure 4m, sleep is increased following training compared with non-trained siblings consistent with previous reports [51, 58]. *R23E10* neurons show higher GFP signal in trained animals when assessed the following morning compared with their naïve counterparts indicating that sleep-promoting neurons are more active following courtship memory training (Figures 4n and 4o for quantification). Together these data indicate that the response properties of *R23E10* neurons display long-lasting changes following protocols that induce LTM.

### Prior feeding experience alters the recruitment of Dopamine receptors to modulate sleep

Initial studies indicated that the wake promoting effects of Dopamine on dFB neurons are mediated by the *Dopamine 1-like receptor 1* (*Dop1R1*) [35, 36, 61]. However, the role of the *Dop1R1* has been called into question [19]. We have recently shown that the constellation of receptors expressed on a neuron can be altered by starvation to include a receptor that is not typically present [3]. Specifically, our data suggest that the new receptor is recruited to amplify wake-promoting signals in clock neurons to allow animals to engage in adaptive waking behaviors. To determine whether the phenomenon of recruiting new wake-promoting receptors to a neuron will generalize to the dFB, we re-examined the role of the *Dop1R1* in *R23E10* neurons. In addition, we also evaluated *R55B01* neurons because these cells have a limited response to Dopamine. We hypothesized that if the *Dop1R1* is needed to support waking following a metabolic challenge, knocking it down would manifest as an increase in sleep. Since 18 h of starvation did not change the response properties of *R23E10* neurons (Figures 4g,h), we used time-restricted feeding to safely impose a greater and more sustained metabolic challenge to the fly [62–64]. That is, time-restricted feeding should provide a substantial metabolic challenge but should not result in lethality. The time-restricted feeding protocol is shown in Figure 5a. Flies are only given access to food between 8am and 5pm for a total of 7 days (restricted). After time restricted feeding, flies are placed into Trikinetics tubes where they were allowed to eat *ad lib* while sleep is evaluated. Siblings that were maintained in vials with standard food available *ad lib* and flipped at the same times as their restricted counterparts served as treatment controls (Figure 5a). Consistent with previous reports [19, 36], knockdown of *Dop1R1* in *R23E10* neurons did not alter sleep in flies that were able to feed *ad lib* (Figures 5b,d,e). Similarly, no changes in sleep were observed in untreated *R55B01>Dop1R1^RNAi^* flies (Figures 5c,d,e). However, both *R23E10>Dop1R1^RNAi^* and *55B01>Dop1R1^RNAi^* flies displayed dramatic increases in sleep following 7 days of time-restricted feeding compared to their *R23E10/+, 55B01/+* and *Dop1R1^RNAi^/+* parental controls (Figures 5f-i). These data are consistent with the hypothesis that, under certain circumstances, a new receptor can be recruited to amplify wake-promoting signals.

**Figure 5.**
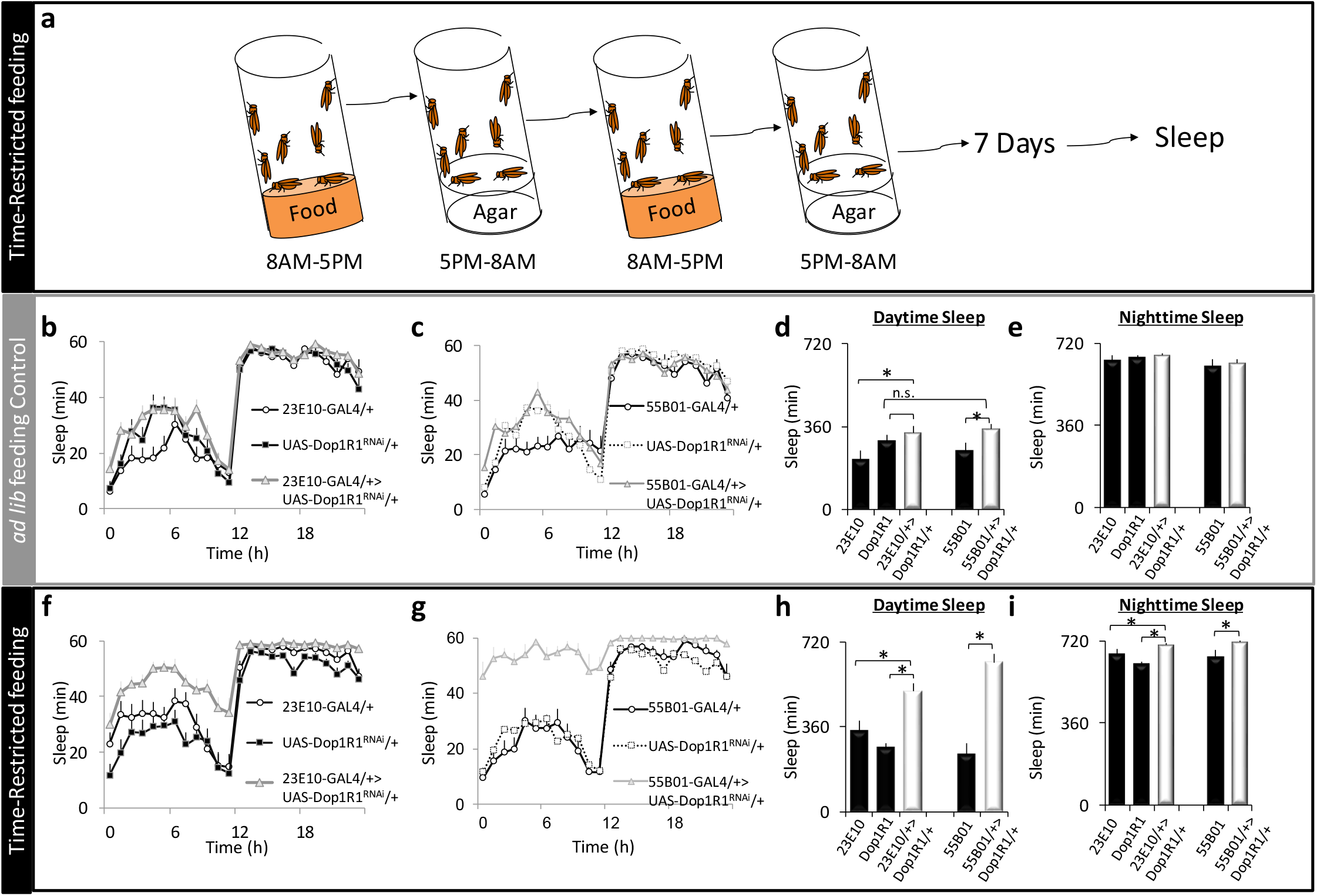
Time-restricted feeding increases sleep via the Dop1R1. **a)** Protocol for time-restricted feeding **b,c)** During *ad lib* feeding, sleep in *R23E10>Dop1R1^RNAi^* and *R55B01>Dop1R1^RNAi^* flies does not differ from *R23E10/+, Dop1R1^RNAi^/+* or *R55B01/+* parental controls (n=13-16 flies/group ANOVA for Genotype F[2,42]=2.8, p=0.07 and ANOVA for Genotype F[2,41]=1.9, p=0.15 for R23E10 and R55B01 respectively. **d)** Daytime sleep was not altered in either *R23E10>Dop1R1^RNAi^* or *R55B01>Dop1R1^RNAi^* flies compared to both parental controls (ANOVA for Genotype F[2,42]=2.8, p=0.02 and F[2,41]=2.9, p=0.06, *p<0.05 Modified Bonferroni Test **e)** Nighttime sleep was not changed in either *R23E10>Dop1R1^RNAi^* or *R55B01>Dop1R1^RNAi^* flies compared to parental controls (ANOVA for Genotype F[2,42]=0.9, p=0.40 and ANOVA for Genotype F[2,41]=0.79, p=0.46, *p<0.05 Modified Bonferoni Test. **f,g)** Following time-restricted feeding, sleep is increased in *R23E10>Dop1R1^RNAi^* and *R55B01>Dop1R1^RNAi^* flies compared to *R23E10/+, Dop1R1^RNAi^/+* or *R55B01/+* parental controls (n=13-16 flies/group ANOVA for Genotype F[2,40]=68.8, p=9.9^eE-16^ and ANOVA for Genotype F[2,35]=38.1.8, p=8.1^eE-10^ for R23E10 and R55B01 respectively. **h)** Daytime sleep was significantly increased in *R23E10>Dop1R1^RNAi^* and *R55B01>Dop1R1^RNAi^* flies following time restricted feeding (ANOVA for Genotype F[2,40]=13.6, p=2.96^E-05^ and F[2,35]=30.5, p=2.07^E-08^, *p<0.05 Modified Bonferroni Test **i)** Nighttime sleep was not changed in either *R23E10>Dop1R1^RNAi^* or *R55B01>Dop1R1^RNAi^* flies compared to parental controls (ANOVA for Genotype F[2,40]=11.41, p=0. 0.0001 and ANOVA for Genotype F[2,35]=8.9, p=0.0007, *p<0.05 Modified Bonferoni Test. Error bars represent SEM

Time restricted feeding is characterized by defined intervals of feeding and fasting. Thus, during time-restricted feeding, the *Dop1R1* may be recruited to *R23E10* neurons to either support waking during starvation or to support waking when food is available. To distinguish between these possibilities, we monitored sleep in *R23E10>Dop1R1^RNAi^* and *55B01>Dop1R1^RNAi^* flies and their parental controls during baseline, during 18 h of starvation and for three days after being placed back onto food (Figures 6a,b). As above, no changes in baseline sleep were observed while knocking down *Dop1R1* in either *R23E10* or *R55B01* neurons compared to parental controls (Figures 6a,b,c,d). Moreover, *R23E10>Dop1R1^RNAi^* and *55B01>Dop1R1^RNAi^* flies exhibited similar sleep patterns to parental controls during starvation (Figure S8a). However, during recovery following 18h of starvation, an increase in sleep was observed in both *R23E10>Dop1R1^RNAi^* and *55B01>Dop1R1^RNAi^* flies compared to their respective parental controls (Figures 6a,b). It is important to highlight that sleep in the experimental lines only diverged from their parental controls after several hours, or more, of recovery (Figures 6a,b arrows). Quantification of sleep during Recovery day 1 and Recovery day 2 is shown in Figures S8b,c. Sleep stabilized in both *R23E10>Dop1R1^RNAi^* and *55B01>Dop1R1^RNAi^* flies on the third day of recovery and remained elevated for several days thereafter (Figures 6,a,b,e and data not shown). Thus, these data indicate that the *Dop1R1* is recruited to *R23E10* and *R55B01* neurons to support waking behavior during recovery from starvation.

**Figure 6.**
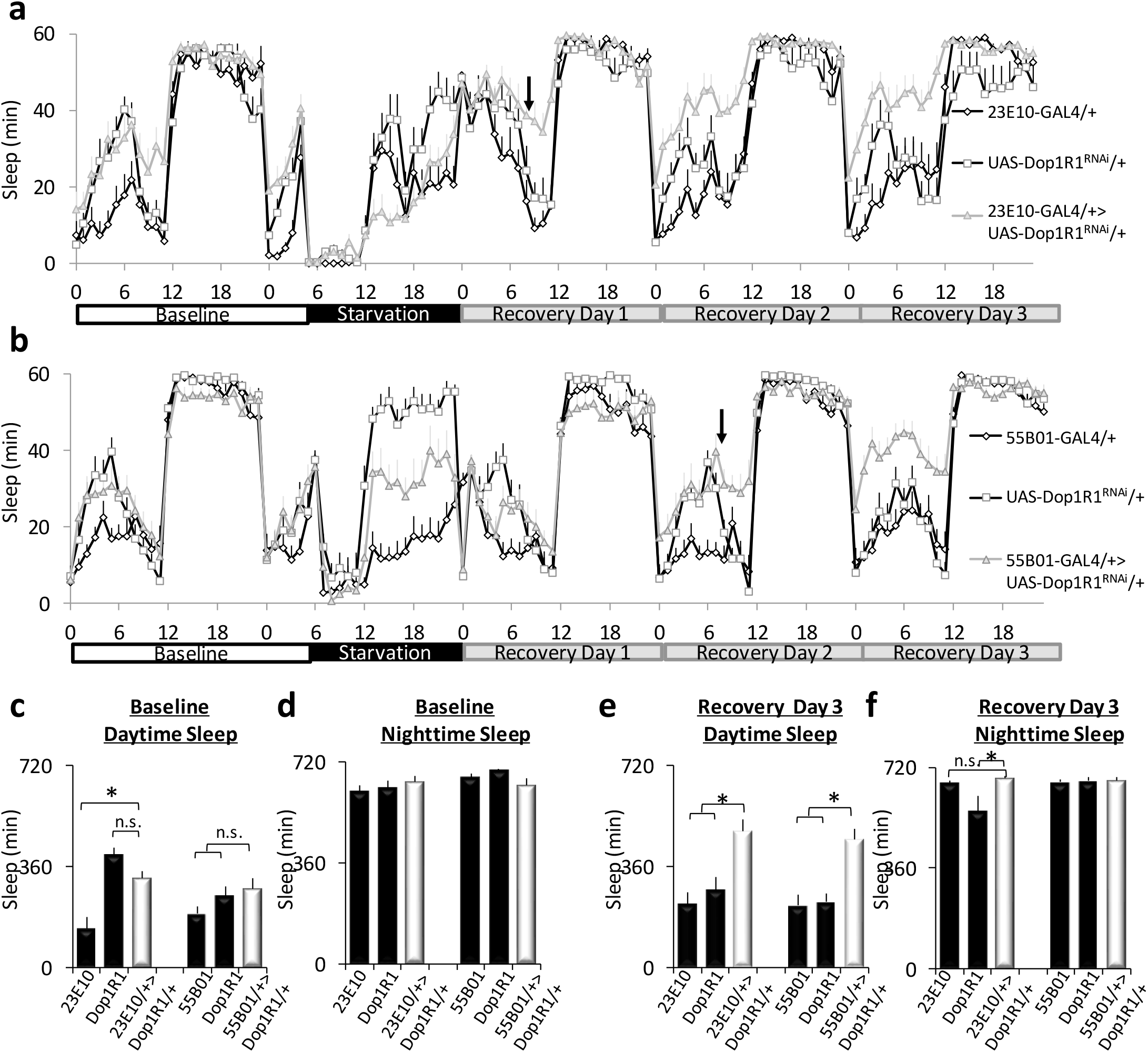
Starvation alters recovery sleep and the response properties of sleep promoting neurons. **a-b)** Sleep profiles in *R23E10>Dop1R1^RNAi^*, *R55B01*>*Dop1R1^RNAi^* flies and their parental controls *R23E10/+, Dop1R1^RNAi^/+* and *R55B01/+* during baseline, 18 h of starvation and three days of recovery (n=13-16 flies/group). **c, d)** During baseline, no changes in daytime or nighttime sleep were observed in *R23E10>Dop1R1^RNAi^* or *R55B01>Dop1R1^RNAi^* flies compared to both parental controls (Daytime: ANOVA F[2,36]=8.1, p=0.001 and F[2,38]=2.26, p=0.11), (Nighttime: ANOVA F[2,36]=1.5, p=0.22 and ANOVA F[2,38]=1.6, p=0.20, *p<0.05 Modified Bonferoni Test. **e)** On recovery Day 3, daytime sleep is increased in *R23E10>Dop1R1^RNAi^* and *R55B01>Dop1R1^RNAi^* flies compared to *R23E10/+, Dop1R1^RNAi^/+* or *R55B01/+* parental controls (ANOVA F[2,36]=9.1, p=0.0006 and F[2,38]=12.1, p=8.43^E-05^. **f)** Nighttime sleep is not changed in *R23E10>Dop1R1^RNAi^* and *R55B01>Dop1R1^RNAi^* flies compared to *R23E10/+, Dop1R1^RNAi^/+* or *R55B01/+* parental controls (ANOVA F[2,36]=3.9, p=0.02 and F[2,38]=0.09, p=0.9 *p<0.05 Modified Bonferoni Test. Error bars represent SEM

To gain additional insight into the physiological impact of starvation, we used live-brain imaging to evaluate the responses of *R23E10>Epac; UAS-Dop1R2^RNAi^* and *R55B01>Epac; UAS-Dop1R2^RNAi^* to Dopamine under baseline and on recovery day 2 from starvation. While both *Dop1R1* and *Dop1R2* couple to *Gαs, Dop1R2* appears to also activate *Gαq* [65]. We hypothesized that knocking down *Dop1R2* would reduce the response of *R23E10* neurons to dopamine under baseline conditions and that the neurons would respond to Dopamine following 2 days of recovery from starvation. *R23E10>UAS-Dop1R2^RNAi^* flies displayed an increase in sleep under baseline conditions consistent with previous reports (data not shown)[19]. As seen in Figures 7a,b *R23E10>Epac; UAS-Dop1R2^RNAi^* neurons did not show a strong response to Dopamine during baseline. However, during recovery from starvation *R23E10>Epac; UAS-Dop1R2^RNAi^* neurons exhibited a significant increase in their response to Dopamine suggesting a new receptor is added (Figures 7a,b). Next we evaluated the effects of starvation on *R55B01* neurons. As seen in Figures 7c,d neither *R55B01>Epac* nor *R55B01>Epac;UAS-Dop1R2^RNAi^* responded to Dopamine under baseline conditions. Although much smaller than that observed for *R23E10* neurons, *R55B01>Epac;UAS-Dop1R2^RNAi^* neurons did respond modestly to Dopamine after two days of recovery from starvation (Figures 7c,d). While we cannot exclude a role of other Dopamine receptors (e.g. Dopamine/Ecdysteroid receptor) for the observed changes, when viewed with the sleep experiments shown above (Figures 5 and 6), these data suggest that during recovery from starvation the constellation of Dopamine receptors in *R23E10* and *R55B01* changes and most likely includes the recruitment of *Dop1R1*. These data provide new insights into the mechanisms used by sleep-circuits to link internal states and prior waking history with sleep need.

**Figure 7.**
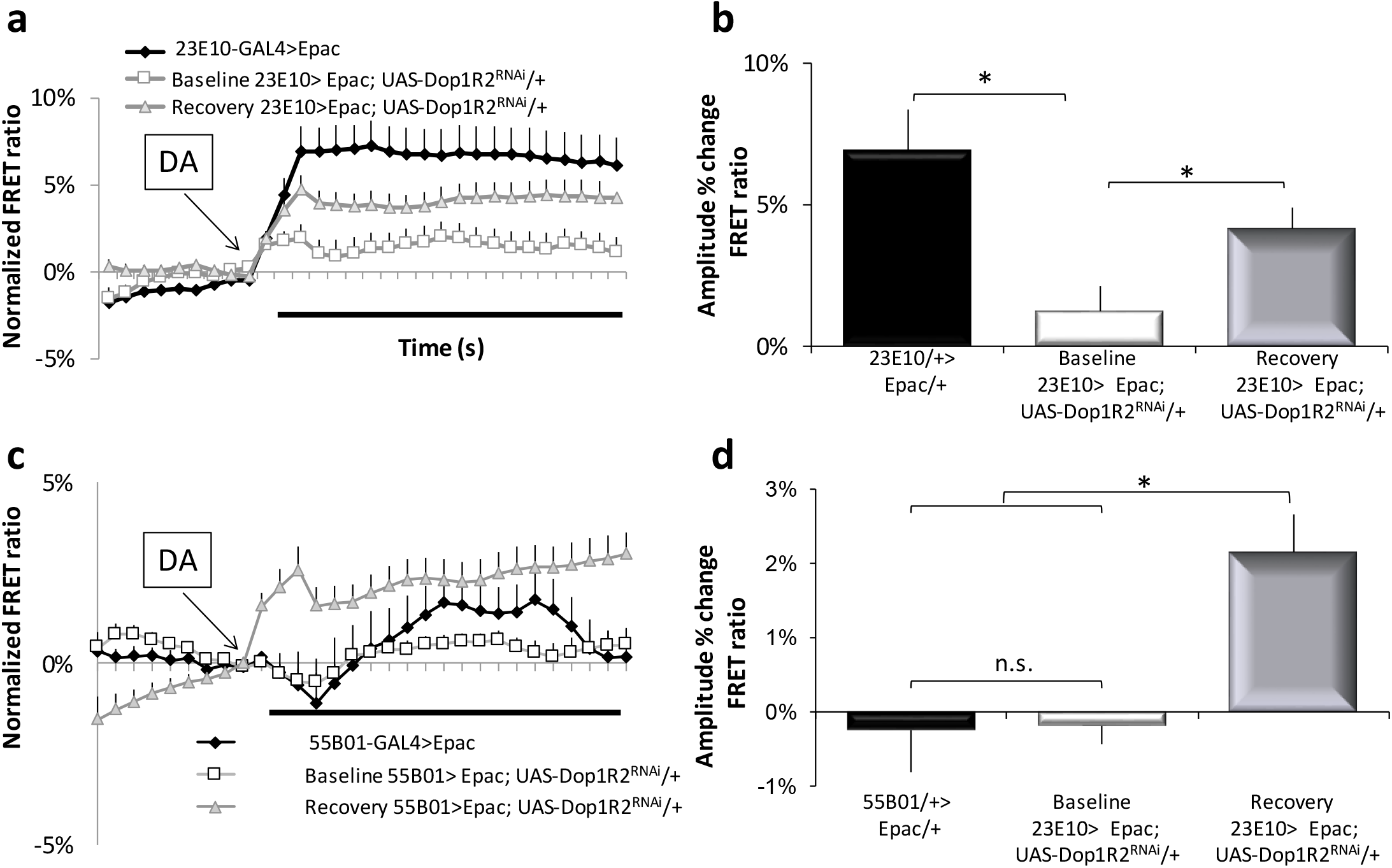
Starvation alters the response properties of sleep promoting neurons. **a)** R23E10 cells expressing the cAMP sensor Epac1-camps respond to 3 × 10-5 M DA (black line; n=22 neurons); the response is dramatically reduced in *R23E10>Epac; Dop1R2^RNAi^* (white line; n = 25 neurons) and partially restored during recovery from 18 h of starvation (grey line; n =29 neurons). **b)** Quantification of a; F[2,73]=7.7, p=0.0009 *p<0.05 Modified Bonferoni Test. **c)** The response of *R55B01>Epac* (black; n =13) and *R55B01>Epac; Dop1R2^RNAi^* (white; n=19) neurons to dopamine is very low; following recovery from starvation, the response of R55B01 neurons to DA increased (grey line; n =20). **d)** Quantification of c; F[2,49]=9.9, p=0.0002 *p<0.05 Modified Bonferoni Test. Error bars represent SEM

## Discussion

In this work we ask whether the activity of sleep-promoting dFB neurons reflects the summation of their upstream inputs in a winner-take all strategy, or if they can change their own response properties to influence how environmental demands alter behavioral state. Although one possibility does not preclude the other, our data indicate that both time-restricted feeding and ~18 h of starvation, alter the constellation of Dopamine receptors expressed by *R23E10* and *R55B01* neurons. These results emphasize that the ability of upstream circuits to alter behavior will not only depend upon the strength of the incoming signals but also the recent historical context of *R23E10* and *R55B01* neurons themselves. In that regard, it important to note that while sleep competes with all motivated waking-behavior, the proper regulation of motivated behaviors degrade in the absence of sleep [50, 66–74]. Together these observations suggest that it will not be possible to fully understand sleep’s function without knowing how sleep-promoting neurons alter their functional properties to prioritize conflicting motivational states.

### The dFB is comprised of independent sets of sleep promoting neurons

Previous studies indicate that the dFB can independently gate different aspects of sleep [26].

Moreover, the expression of the gap junction *innexin6* in *R23E10* neurons is required to gate sensory thresholds and sleep time[48]. Together these data suggest the presence of independent sets of dFB neurons. Although we screened through a refined set of hand selected GAL4 lines that project into the dFB in a manner similar to *104y-GAL4* and *C5-GAL4* [10, 49], we only identified one additional GAL4 line, *R55B01*, that reliably increased sleep when crossed with *UAS-dTrpA1*. Similar results were obtained when activating the neurons using the bacterial sodium channel *UAS-NaChBac*. This latter result is relevant given that sleep is modulated by temperature and that GABAergic neurons projecting onto dFB regulate temperature-dependent changes in sleep [75, 76]. We also identified dFB projecting GAL4 lines that increased waking. However, the precise role that these GAL4 lines play in waking awaits additional inquiry. Nonetheless, these data suggest that sleep promoting dFB neurons are limited in number. It should be noted that the extent of the overlap between *R23E10* and *R55B01* remains unclear as the *R23E10-LexA* driver does not fully recapitulate the *R23E10-GAL4* expression patterns (Figure S5). Although it may be possible to subdivide *R23E10* and *R55B01* neurons further, our data suggest that there are at least two independent sets of sleep promoting neurons that have projections into the dFB (see below for additional discussion). In mice, sleep promoting neurons are also diverse as assessed by their role in regulating behavior and their projection patterns [77].

### Allatostatin inhibits sleep-promoting neurons

Although AstA-expressing neurons are believed to promote sleep by releasing glutamate onto *R23E10* neurons [25], our data identify a wake-promoting effect of AstA on *R23E10* neurons. Specifically, we find that knocking down either the *AstAR1* or *AstAR2* in *R23E10* or *R55B01* neurons using several independent RNAi lines results in a substantial increase in sleep. The observation that knocking down either the *AstAR1* or *AstAR2* in *R23E10* neurons increases sleep, suggests that *R23E10* neurons are under constant inhibitory tone from AstA and that both receptors play an important role in modulating sleep in untreated flies. In addition, we report that the application of 1mM ATP to *R23E10>Epac; AstA-GAL4>P2X2* neurons reduces cAMP levels similarly to that observed with application of AstA. Although our data appear to be in conflict with that of Ni and colleagues, we hypothesize that the co-release of inhibitory AstA with excitatory glutamate may attenuate the overexcitement of *R23E10* neurons during high sleep drive and allow animals to maintain wakefulness in dangerous or life-threatening conditions.

In support of this hypothesis, a growing body of evidence indicates that sleep and wake regulating neurons co-release different transmitters and neuropeptides to better match sleep-need with environmental demands [78]. Indeed, it has been suggested that antagonistic neurotransmitter co-release may both prevent excessive excitation of postsynaptic targets and increase the flexibility of neuronal networks [45, 78, 79]. For example, the co-release of GABA by histaminergic *tuberomammillary nucleus* is believed to prevent histamine-induced overexcitement of downstream circuits and thereby better support normal amounts of waking [80]. Similarly, the co-release of galanin by noradrenergic neurons is believed to prevent overexcitement of *locus coeruleus[81]*. Although the co-expression of AstA and glutamate has been established [25, 82], our data reveal a new mechanism for how incompatible motivational drives can be regulated.

An alternate possibility to explain the wake-promoting effects of AstA on *R23E10* and *55B01* neurons may be that *R23E10* and *R55B01* are, in fact, a heterogeneous sets of neurons which also include wake promoting neurons and/or neurons whose primary role is to promote active sleep at the expense of deep sleep [83]. Active sleep is best observed following 24 h of sleep deprivation and is associated with dramatic reductions in the response to external sensory stimuli. If AstA were to inhibit a subset of *R23E10* that promote active sleep, one would predict an increased deep sleep. The diversity in the responses to Dopamine seen in both *R23E10* and *R55B01* neurons (Figure 3f) further supports the hypothesis that these GAL4 lines express in heterogeneous set of neurons with different functions; a previous study has found similar heterogeneity in *R23E10* neurons. [38]. Alternatively, it is possible that AstA does not inhibit the *R23E10* neurons *per se* but rather alters temporal spike patterns to favor active-sleep or deep-sleep in a manner similar to that observed in clock neurons [84]. Indeed, AstA has been shown to alter the spike-timing precision of mechanoreceptor afferents in *Carcinus maenas* and the mammalian homologue of AstA, *Galanin*, alters spontaneous spike firing in rodents [85, 86]. These possibilities will be explored in future studies.

It is important to note that neuropeptides are notoriously pleiotropic and also work as neurohormones [44]. Thus, it is possible that additional sleep and wake promoting circuits downstream of *AstA-GAL4* or *65D05-GAL4* will be found elsewhere in the brain. It is interesting to note that neurons expressing the mouse homologue of AstA, galanin, also regulate conflicting behaviors [77]. Specifically, Galanin expressing neurons that project to the *tuberomammillary nucleus* promote sleep while Galanin expressing neurons that projection to the medial amygdala promote parental behaviors [77, 87, 88]. Thus it will be important for future studies to discern how AstA and galanin circuits regulate competing activities in other circuits.

### R23E10 neurons differentially encode arousal signals

Increased sleep drive may be maladaptive in many circumstances since falling asleep could place the individual in danger of immediate physical harm [7, 8]. Increased sleep also competes with important waking behaviors such as foraging, eating and mating [3, 4]. However, sleep plays a critical role in learning and memory, supports adaptive behavior and facilitates creative insight [9–12]. How do sleep promoting neurons distinguish between competing drives? Our live-brain imaging data suggest the intriguing possibility that *R23E10* neurons can distinguish between different types of waking and change their response properties accordingly. Specifically, *R23E10* neurons selectively <u>decrease</u> their response to the wake-promoting effects of Dopamine following conditions that promote plasticity. In contrast, *R23E10* neurons <u>increase</u> their response to the wake-promoting effects of AstA during conditions when sleep drive is high but the expression of sleep might be dangerous (e.g. during sleep deprivation). Finally, *R23E10* neurons appear to contain a memory trace of starvation as evidenced by the increase inhibitory tone conveyed by the recruitment of the *Dop1R1*. Thus, *R23E10* neurons are plastic and can utilize AstA and DA in very different ways to favor specific behavioral outcomes.

As mentioned, increasing inhibitory signals onto sleep promoting neurons during sleep deprivation may be an adaptive response that allows animals to stay awake despite increasing sleep drive. Indeed, lesioning Galanin neurons in the preoptic area reduces sleep rebound following sleep loss in mice [89]. However, AstA is also reported to be a satiety signal that limits feeding [27, 32, 37, 90]. The effects of AstA on feeding seem to be in conflict with the studies linking sleep deprivation to increased food intake [91–94]. Nonetheless, recent reports have identified complex interactions between hunger and satiety signals that may be nutrient specific and can be modified by Hebbian plasticity [32, 34, 40, 44]. Thus, it is reasonable to expect that sleep deprivation must differentially activate neurons regulating a variety of motivated behaviors including satiety and sleep drive. Indeed, these data indicate that sleep deprivation may be harnessed as a tool to further explore how internal states impact neuronal circuits regulating conflicting goal-directed behaviors [30].

### R23E10 neurons support Long Term Memory

Interestingly, the ventral Fan Shaped body (vFB) promotes sleep and regulates the activity of Dopaminergic aSP13 neurons (DAN-aSP13s) to consolidate long-term memory as assessed using courtship conditioning [59]. Surprisingly, activating *R23E10* neurons does not alter the activity of DAN-aSP13s neurons suggesting that the dFB may not be involved in courtship memory [59]. It should be noted, however, that *R23E10* neurons regulate the activity of other sets of arousal promoting neurons including the MV1 Dopaminergic neurons and octopaminergic arousal neurons [25, 95]. These data suggest that the dFB and vFB may play distinct roles in different memory assays. Nonetheless, our data clearly indicate that *R23E10* neurons change their response properties following training that induces LTM but not massed training. Importantly, the changes in *R23E10* neurons only return to baseline 48h after training. It is possible that *R23E10* neurons play an important, yet indirect, role in courtship memory. That is, activating *R23E10* neurons strongly suppresses the response to external sensory stimuli [48, 83]. Thus the activity of *R23E10* neurons may protect sleep by limiting the opportunity for external stimuli to wake the animal up. Under this scenario, dFB and vFB neurons would work in concert to carry out sleep functions.

In addition, we find that courtship conditioning increases the activity of *R23E10* neurons as assessed by *CaLexA*. Consistent with this observation, spaced-training reduced the inhibitory effects of Dopamine on *R23E10* neurons. Importantly massed-training, which only induced short-term memory, does not alter the response of *R23E10* neurons to Dopamine. A previous report has shown that the ectopic expression of *Dop1R1* in dFB neurons reduces sleep and fails to rescue courtship conditioning memory in *Dopamine transporter* mutants (*fmn*) [36]. Together these data emphasize that reducing Dopaminergic signaling to the dFB neurons is important for long-term memory following courtship conditioning.

### Internal state remodels dopamine receptors in sleep promoting neurons

The arousal promoting properties of Dopamine projections to the dFB are well established [19, 25, 35, 36, 61]. However, which Dopamine receptor is responsible for the increased waking remains controversial. Initial results, using the ectopic expression *Dop1R1* and live-brain imaging implicated the *Dop1R1* [35, 36]. Nonetheless, RNAi knockdown of *Dop1R1* in dFB neurons did not alter sleep [35]. A subsequent study demonstrated that activation of the *Dop1R2* results in a transient hyperpolarization of dFB neurons followed by a lasting suppression of excitability lasting minutes; knocking down the *Dop1R2* but not *Dop1R1* altered sleep [19]. Our data replicate previous RNAi studies that have failed to observe a change in sleep following the expression of *Dop1R1^RNAi^* in dFB neurons in untreated flies. Indeed, no changes in FRET signal were observed upon the bath application of Dopamine to untreated *R23E10>UAS-EPAC; Dop1R2^RNAi^* or *R55B01>UAS-EPAC; Dop1R2^RNAi^* flies further supporting the role of *Dop1R2* in baseline sleep regulation.

However, in the context of time-restricted-feeding and starvation, knocking down the *Dop1R1* in *R23E10* and *R55B01* neurons resulted in substantial increases in sleep. We hypothesize that both time-restricted feeding and starvation place energy saving sleep-drive in conflict with a metabolic signal that facilitate motivated waking behavior. Increased wake-drive would allow the animals to be more alert and productive during their primary wake-period. Under these circumstances, the *Dop1R1* would be recruited to provide additional inhibitory tone to sleep promoting *R23E10* and *R55B01* neurons during the light period. Interestingly, starvation results in the recruitment of the *Pigment dispersing factor receptor* (*Pdfr*) into wake-promoting *large ventrolateral neurons* (*lLNvs*) to facilitate waking at the end of the biological day [3]. Whether other sleep and wake-promoting circuits are regulated in a similar fashion is an open question that will require additional inquiry.

## Conclusions

Given the important roles that Dopamine and AstA play in regulating motivated behaviors, including sleep, we studied their impact on a subset of sleep promoting neurons. Our data reveal that the co-release of AstA is wake promoting and likely serves to maintain waking during periods of high sleep drive. In addition, our data reveal that time-restricted feeding and 18 h of starvation recruits the *Dop1R1* to sleep promoting neurons to maintain wakefulness during the day. These results are consistent with our previous data showing that the wake-promoting lLNvs recruit the *Pdfr* following sleep loss to facilitate waking and that wing-cut reactivates a developmental sleep circuit [3, 96]. The ability of sleep and wake promoting neurons to alter their own physiology, including the recruitment of a new receptor, provides important clues into sleep regulation and function.

## Methods

### Flies

Flies were cultured at 25°C with 50-60% relative humidity and kept on a diet of yeast, dark corn syrup and agar under a 12-hour light:12-hour dark cycle. *R23E10-GAL4; R55B01-GAL4; R33E06-GAL4; R19C06-GAL4; R28H10-GAL4; R09D11-GAL4; R26B11-GAL4; R84C10-GAL4; R58G11-GAL4; R24E05-GAL4; R65C03-GAL4;R12D12-GAL4* and *LexAop-TrpA1* were a kind gift from Dr Gerald Rubin at Janelia Farms Research campus. *AstA-GAL4; R65D05-LexA; R23E10-LexA; UAS-Dcr2; UAS-TrpA1; UAS-GFP; UAS-RFP, LexAop-GFP; AstA-R1^RNAiJF02578^; AstA-R2^RNAiJF01955^*; *Goα47A^RNAi HMS01129^; Giα65A^RNAi GL00328^; Giα65A^RNAi HMS01273^; Gβ5^RNAi JF02941^; Gβ13F^RNAi JF01609^; Gβ76C^RNAi JF03127^; Gγ1^RNAi HMS01361^; UAS-Dop1R1^RNAi62193^; UAS-Dop1R2^RNAi51423^; UAS-FMRFaR ^RNAi25858^; UAS-Dh44-R2 ^RNAi29610^; UAS-MsR1^RNAi27529^; UAS-Dh44-R1^RNAi28780^: UAS-, cchamideR^RNiI51168^; UAS-NepyR^RNAi25944^; UAS-CapaR^RNAi27275^ UAS-Pk2R1^RNAi 29624^; UAS-CCKLR-17D1^RNAi67865^; UAS-DH31R^RNAi25925^: UAS-AstC-R1^RNAi27506^; UAS-TkR99D^RNAi 27513^: UAS-MsR2^RNAi25832^; UAS-Lkr^RNAi25936^; UAS-CrzR^RNAi26017^; UAS-ProcR^RNAi29414^*were obtained from the Bloomington stock center. *UAS-Epac1.camps* was obtained from Paul Taghert (Washington University in St. Louis). *UAS-P2X2* and *LexAop-Epac1.camps* were a kind gift from Orie Shafer (City University of New York). *AstA-R1^RNAiv39221^* and *AstA-R1^RNAiv3400^* were obtained from the Vienna *Drosophila* RNAi stock center (Austria). *AstA-R1^RNAiHMJ21471^* was obtained from the Kyoto *Drosophila* stock center (Japan). *CaLexA* flies were obtained from Jin Wang (UCSD).

### Sleep

Sleep was assessed as previously described [52]. Briefly, flies were placed into individual 65 mm tubes containing the same food as they were reared on. All activity was continuously measured through the Trikinetics Drosophila Activity Monitoring System (www.Trikinetics.com, Waltham, Ma). Locomotor activity was measured in 1-minute bins and sleep was defined as periods of quiescence lasting at least 5 minutes. All sleep experiments were replicated a minimum of two times.

### Sleep Deprivation

Sleep deprivation was performed as previously described [50, 73]. Briefly, flies were placed into individual 65 mm tubes and the sleep-nullifying apparatus (SNAP) was used to sleep deprive these flies for 12 hours during the dark phase (lights out to lights on).

### Starvation, Time-Restricted Feeding

For starvation, flies were individually placed into Trikinetics tubes containing the same food they were reared on and sleep was monitored for 2 days. On the morning of day 3, flies were placed into tubes containing 1% agar and monitored for 18 h. the flies were then placed back into Trikinetics tubes with food. For time-restricted feeding, flies were housed in vials on standard food between 8am and 5pm. At 5pm, flies were transferred to vials containing 1% agar until 8am the next morning when they were transferred to a new vial containing food. Flies underwent this protocol for 7 days. After 7 days, flies were individually placed into Trikinetics tubes containing standard food where they were allowed to eat *ad lib*. Siblings that were maintained in vials with standard food available *ad lib* and flipped at the same times as their restricted counterparts served as treatment controls

### Social Enrichment

Social enrichment was performed as previously described [51]. Briefly, flies were housed in groups of 20 until 3-4 day old. Then, flies were separated in isolated or enriched groups. Isolated flies were transferred in individual 65 mm tubes. Enriched flies were pooled in groups of 50 flies for 5 days.

### Courtship Conditioning

Training for naïve males was based on previously described methods [10]. For LTM, a spaced training protocol consisting of three 1-hour training sessions with a mated female, each separated by one hour was employed. For massed training protocol that does not induce LTM, a single 3h pairing with a mated female was employed.

### Live Brain Imaging

Flies were chilled for approximately 5 minutes prior to pinning them onto a sylgaard dissection dish. Brains were dissected in calcium-free HL3 and then transferred onto a poly-lysine treated dish (35 3 10 mm Falcon polystyrene) containing 3 ml of 1.5mM calcium HL3. Two to four brains were assayed concurrently, typically untreated controls vs a comparison group (e.g. sleep deprived, starved, etc). Image capture was done using an Olympus BX61 and x,y,z stage movements were set via SLIDEBOOK 5.0 (Intelligent Imaging Innovations), which controlled a Prior H105Plan Power Stage through a Prior ProScanII. Multiple YFP/CFP ratio measurements were recorded in sequence from each brain in the dish. Following baseline measurements, 1 ml of saline containing either AstA or DA, was added to the bath (dilution factor of 1/4). We used synthetic AstA (Neo-MPS) and dopamine (Sigma-Aldrich). Crustacean cardioactive peptide (CCAP), drosophila myosuppressin (DMS), allatostatin C (astA C), proctolin, TPAEDFMRFamide, corticotropin-releasing factor-like diuretic hormone 44 (DH44), Tachykinin 1, Tachykinin 3, short neuropeptide F (sNPF), adipokinetic hormone (AKH), corazonin, and melatonin were kindly gifted by Paul Taghert (Washington University in St. Louis). For further details see [42, 43]. For experiment using P2X2, 1 mM ATP was used to activate the P2X2 receptor.

### Immunocytochemistry

Flies were fixed in 4% PFA, brains were dissected in ice cold PBS and incubated overnight with the following primary antibodies: mouse anti-AstA, (5F10, 1:2 dilution, Hybridoma Bank, University of Iowa), chicken anti-GFP (GFP-1020; 1:1000, Aves Lab), rabbit anti-dsRed (Living Colors DsRed Polyclonal Antibody, 1:250, Clontech). Secondary antibodies were Alexa 488, 594 and 633 conjugated at a dilution 1:200. Brains were mounted on polylysine treated slides in Vectashield H-1000 mounting medium. Confocal stacks were acquired with a 0.5μm slice thickness using an Olympus FV1200 laser scanning confocal microscope and processed using ImageJ.

### Statistics

All comparisons were done using a Student’s T-test or, if appropriate, ANOVA and subsequent planned comparisons using modified Bonferroni test unless otherwise stated. Note that a significant omnibus-F is not a requirement for conducting planned comparisons [97]. All statistically different groups are defined as *P < 0.05.

## Author Contributions

S.D., M.K.K., B.V.S. and P.J.S. designed the experiments and wrote the paper. S.D., M.K.K., L.C., and P.J.S. performed the experiments. S.D., M.K.K., L.C., and P.J.S. analyzed the data. The funders had no role in study design, data collection and analysis, decision to publish, or preparation of the manuscript

## Acknowledgments

We thank Gerald Rubin, Arnim Jenett, Jin Wang, Ori Shafer and Paul Taghert for sharing reagents and flies.

