## Supplemental data for "Sleep promoting neurons remodel their response properties to calibrate sleep drive with environmental demands"

#### Supplemental Figures

**Figure S1**. **Changes in sleep in parental controls following 6h at 31ºC.** % change from baseline sleep at 25ºC seen following switch to 31ºC between 9am and 3pm.

**Figure S2 R23E10 neuropeptide RNAi screen for Daytime Sleep. a)** Individual cAMP responses of *R23E10* neurons expressing UAS-Epac1-camps shown as % change in FRET ratio during exposure to crustacean cardioactive peptide (CCAP), *Drosophila* myosuppressin (DMS), allatostatin C (astA C), proctolin, TPAEDFMRFamide, corticotropin-releasing factor-like diuretic hormone 44 (DH44), Tachykinin 1, Tachykinin 3, short neuropeptide F (sNPF), adipokinetic hormone (AKH), corazonin, and melatonin. **b)** Daytime sleep in female 5-d old flies expressing RNAi lines for the depicted neuropeptide receptors using R23E10 –GAL4 and their parental controls (n=14-16 flies/genotype). To be significant, the experimental lines must be significantly different from both the GAL4/+ (red line) and the UAS/+ (white bar) parental controls : FR: *FMRFamide Receptor*, *DHRRR2*: *Diuretic hormone 44 receptor 2*, *DMSR1*: *Myosuppressin receptor 1*, *DHRRR1*: *Diuretic hormone 44 receptor 1*, *cchamideR*: *CCHamide-1 receptor*, *NepyR*: *RYamide receptor*, *Capar*: *Capability receptor*, *Pk2R1*: *Pyrokinin 2 receptor 1*; *CCKLR*: *Cholecystokinin-like receptor at 17D1A*, *DH31*: *Diuretic hormone 31*,*AstC-R1*: *Allatostatin C receptor 1*, *TkR99D*: *Tachykinin-like receptor at 99D*, *MsR2*: *Myosuppressin receptor 2*, *Lkr*: *Leucokinin receptor*, *CrzR*: *Corazonin receptor*, *Proc*: *Proctolin receptor*. Red line is to facilitate comparisons with R23E10/+ parental control. **c)** RNAi knockdown of the AstA-R1 receptor in *R23E10* neurons did not affect the cAMP response to DA. Error bars represent SEM

**Figure S3 Expressing AstAR1RNAi and AstAR2RNAi R23E10 neurons alters sleep architecture. a, b).** Nighttime sleep bout duration is increased in *UAS-Dcr2, R23E10-GAL4/+> AstA-R1RNAi/+* and *UAS-Dcr2,R23E10-GAL4/+> AstA-R2RNAi/+* experimental flies compared with *UAS-Dcr2, R23E10-GAL4/+, AstA-R1RNAi/+ and AstA-R2RNAi/+* parental controls (n=16/condition, *p<0.05 modified Bonferroni test). **c,d)** Sleep latency is shortened in *UAS-Dcr2, R23E10-GAL4/+> AstA-R1RNAi/+* and *UAS-Dcr2,R23E10-GAL4/+> AstA-R2RNAi/+* experimental flies compared with *UAS-Dcr2, R23E10-GAL4/+, AstA-R1RNAi/+* and *AstA-R2RNAi/+* parental controls (n=16/condition, *p<0.05 modified Bonferroni test). **e)** Total sleep is increased when levels of AstA-R1 is decreased in R23E10 sleep-promoting neurons using 3 additional independent RNAi lines (n=16/condition, *p<0.05 modified Bonferroni test). **f)** Sleep is increased in *AstA-GAL4>UASdTrpA1* and *65D05-LexA>LexAopdTrpA1* flies at 31°C compared with siblings maintained at 25°C; parental controls did not show an increase in sleep at 31°C (n=14-16/condition and genotype, *p<0.05 modified Bonferroni test) Error bars represent SEM.

**Figure S4. Screen of Janelia-GAL4 lines that express in the dorsal Fan Shaped body in a pattern similar to that observed with 104y-GAL4 and C5-GAL4. a)** Despite similar anatomical profiles, most dorsal Fan Shaped Body drivers do not reliably impact sleep when expressing *UAS-Transient receptor potential cation channel A1* and raising the temperature to 31°C (n=14-16 flies/genotype). Results yielded similar results when using UAS-NaChBac (Data not shown). Error bars represent SEM.

**Figure S5. Anatomy of *R23E10* and *R55B01* neurons. a)** Representative confocal stack focusing on the area containing cell bodies of a *R23E10-LexA>LexAop-GFP*, *R55B01-GAL4>UAS-RFP* fly brain stained with anti-GFP antibody**, a')** anti-RFP antibody (magenta) and **a'')** a merged image. Yellow arrows on the merge image indicate cells that express both GFP and RFP. **b)** Quantification of the number of cells expressing only GFP, only RFP or both GFP and RFP. **c)** Representative confocal stack focusing on the area containing the cell bodies of a *R23E10-LexA>LexAop-GFP*, *R23E10-GAL4>UAS-RFP* fly brain stained with anti-GFP antibody **c')** anti-RFP antibody (magenta) and **c'')** a merged image. **d)** Quantification of the number of cells expressing only GFP, only RFP or both GFP and RFP**. e)** Representative confocal stack of a using *55B01-GAL4>UAS-GFP* brain stained with anti-GFP antibody**, e')** anti-AstA antibody (magenta) and **e'')** a merged image. Error bars represent SEM.

**Figure S6. Expressing *AstAR1*$^{RNAi}$ and *AstAR2*$^{RNAi}$ *R55B01* neurons alters sleep architecture.** Sleep parameters in *UAS-Dcr2, R55B01-GAL4/+> AstA-R1*$^{RNAi}$/+, *UAS-Dcr2, R55B01-GAL4/+> AstA-R2*$^{RNAi}$/+ experimental flies and both *UAS-Dcr2*, *R55B01-GAL4/+* and *AstA-R1*$^{RNAi}$/+, *AstA-R2*$^{RNAi}$/+ control flies for**: a,b)** total sleep, **c,d)** Nighttime sleep bout duration, **e,f)** Nighttime sleep latency, **g,h)** Daytime sleep bout duration and **i,j)** counts/waking minute. (n=16/condition, *p<0.05 modified Bonferroni ttest). Error bars represent SEM.

**Figure S7. Quantification of traces shown in Figure 4. a,b)** young (0-1 day old) and adult (6-8 day old) flies. **c,d)** spontaneously short sleeping flies (<400min/day) and normal sleeping siblings (600-900min/day)**. e,f)** Following 12 h Sleep deprivation compared to untreated siblings. **g,h)** Following 18 h starvation compared to untreated siblings. **i,j)** During social enrichment compared to isolated siblings. **k,l)** Following spaced training compared to naïve controls. **m)** Quantification of Figure 4l**,** Space-trained flies vs Naïve controls**.** The reduction of DA responses remained significant 24h after the end of the training but not 48h post-training. A Massed training courtship protocol that does not induce LTM had no significant effect on amplitude of DA response in R23E10 neurons (n=12-25 cells, *p<0.05). Error bars represent SEM.

**Figure S8 Starvation alters recovery sleep**. **a)** During starvation, sleep in *R23E10>Dop1R1*$^{RNAi}$, *R55B01>Dop1R1*$^{RNAi}$ flies is not consistently above or below *R23E10/+*, *Dop1R1*$^{RNAi}$/+ or *R55B01/+* parental controls (n=13-16 flies/group; ANOVA F[2,36]=2.2, p=0.12 and ANOVA F[2,38]=15.4, p=$1.2^{E-05}$, *p<0.05 Modified Bonferoni Test. **b)** Daytime sleep is not increased in *R23E10>Dop1R1*$^{RNAi}$ or *R55B01>Dop1R1*$^{RNAi}$ flies compared to both parental controls *R23E10/+*, *Dop1R1*$^{RNAi}$/+ or *R55B01/+* on Recovery Day 1(ANOVA F[2,36]=4.8, p=0.02 and ANOVA F[2,38]=0.5, p=0.58, *p<0.05 Modified Bonferoni Test. **c)** Sleep is increased in *R23E10>Dop1R1*$^{RNAi}$ and *R55B01>Dop1R1*$^{RNAi}$ flies compared to *R23E10/+*, *Dop1R1*$^{RNAi}$/+ or *R55B01/+* parental controls on Recovery day 2(ANOVA for Genotype F[2,36]=14.25, p=$2.75^{E-05}$ and ANOVA for Genotype F[2,38]=8.35, p=0.0009 for R23E10 and R55B01 respectively.

Figure S1

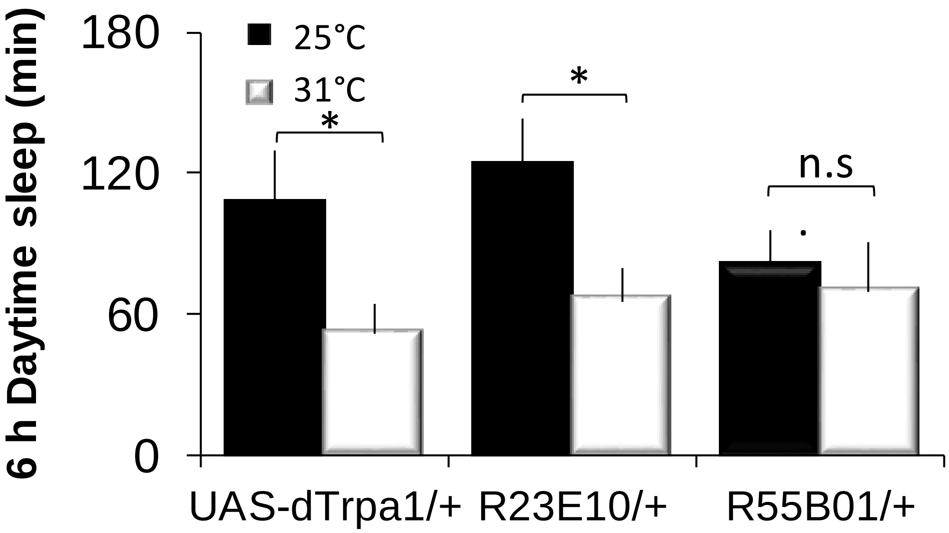

6 h Daytime sleep (min)
180
120
60
0
25℃
31℃
*
*
n.s
UAS-dTrpa1/+
R23E10/+
R55B01/+

Figure S2

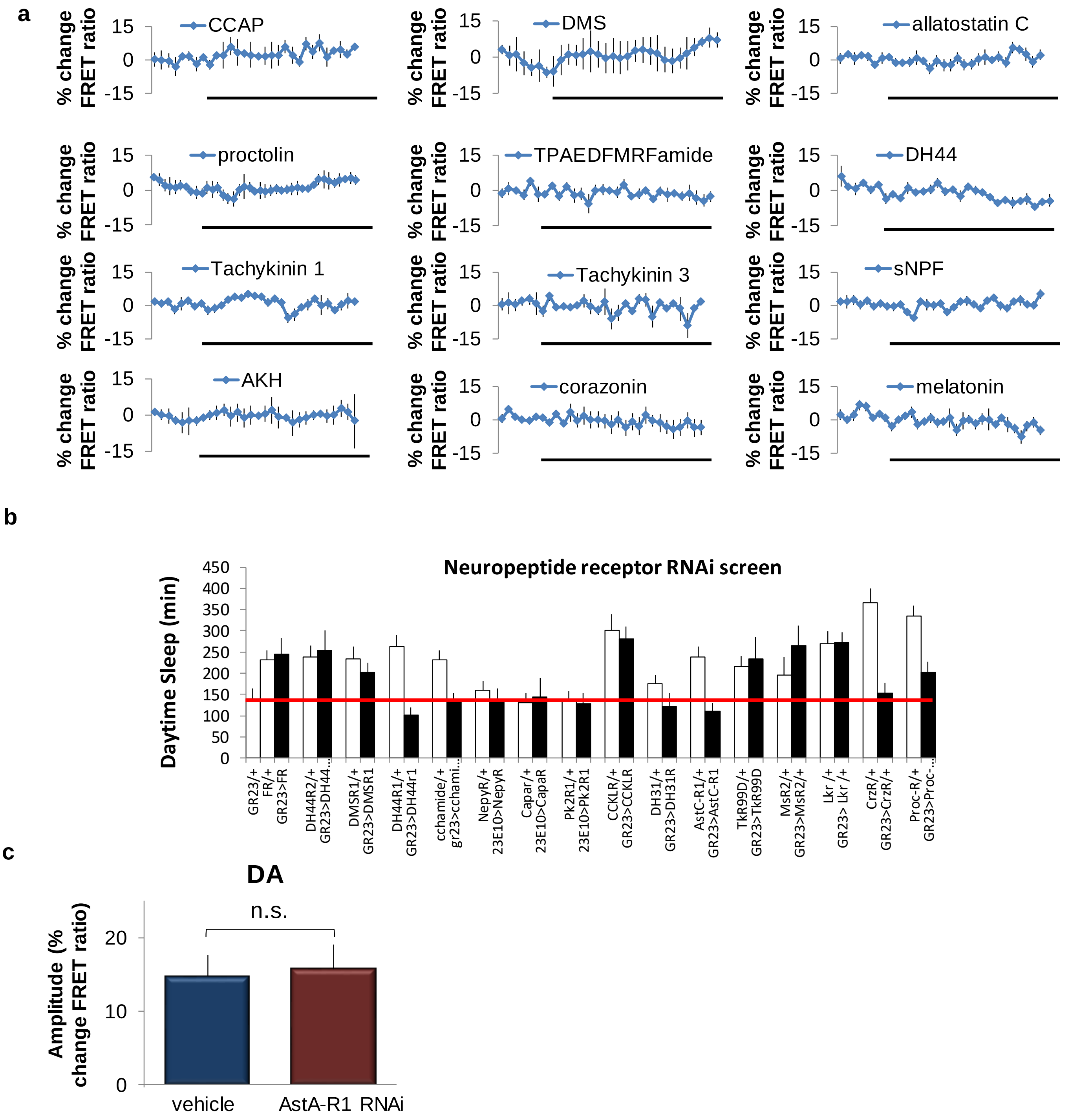

a
% change FRET ratio
15
0
-15
CCAP
DMS
allatostatin C
proctolin
TPAEDFMRFamide
DH44
Tachykinin 1
Tachykinin 3
sNPF
AKH
corazonin
melatonin
b
Neuropeptide receptor RNAi screen
Daytime Sleep (min)
450
400
350
300
250
200
150
100
50
0
GR23/+
FR/+
GR23>FR
DH44R2/+
GR23>DH44...
DMSR1/+
GR23>DMSR1
DH44R1/+
GR23>DH44r1
cchamide/+
gr23>cchami...
NepyR/+
23E10>NepyR
Capar/+
23E10>CapaR
Pk2R1/+
23E10>Pk2R1
CCKLR/+
GR23>CCKLR
DH31/+
GR23>DH31R
AstC-R1/+
GR23>AstC-R1
TkR99D/+
GR23>TkR99D
MsR2/+
GR23>MsR2/+
Lkr /+
GR23> Lkr /+
CrzR/+
GR23>CrzR/+
Proc-R/+
GR23>Proc-...
c
DA
n.s.
Amplitude (% change FRET ratio)
20
10
0
vehicle
AstA-R1 RNAi

Figure S3

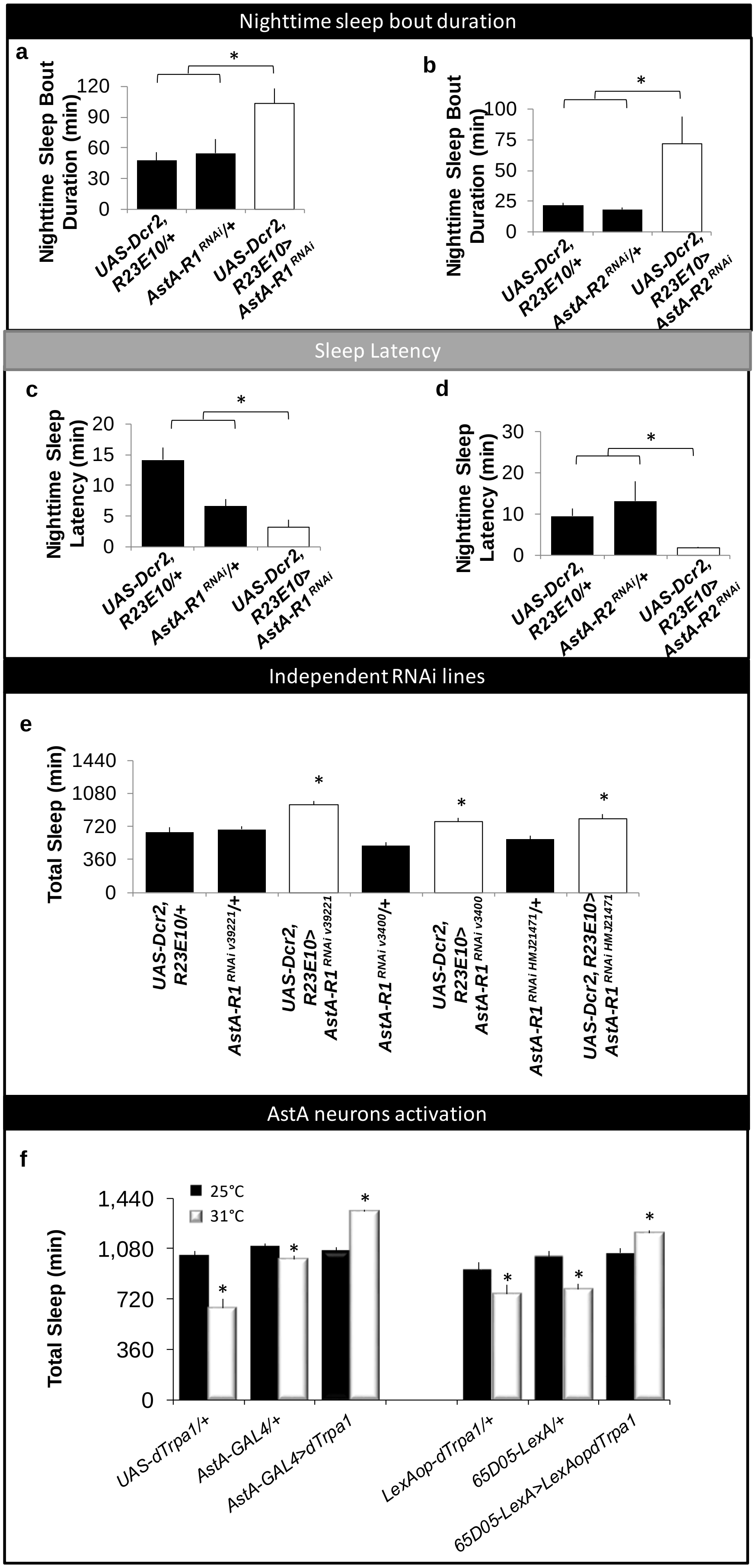

Nighttime sleep bout duration
a
Nighttime Sleep Bout Duration (min)
UAS-Dcr2, R23E10/+
AstA-R1 RNAi/+
UAS-Dcr2, R23E10> AstA-R1 RNAi
b
Nighttime Sleep Bout Duration (min)
UAS-Dcr2, R23E10/+
AstA-R2 RNAi/+
UAS-Dcr2, R23E10> AstA-R2 RNAi
Sleep Latency
c
Nighttime Sleep Latency (min)
UAS-Dcr2, R23E10/+
AstA-R1 RNAi/+
UAS-Dcr2, R23E10> AstA-R1 RNAi
d
Nighttime Sleep Latency (min)
UAS-Dcr2, R23E10/+
AstA-R2 RNAi/+
UAS-Dcr2, R23E10> AstA-R2 RNAi
Independent RNAi lines
e
Total Sleep (min)
UAS-Dcr2, R23E10/+
AstA-R1 RNAi v39221/+
UAS-Dcr2, R23E10> AstA-R1 RNAi v39221
AstA-R1 RNAi v3400/+
UAS-Dcr2, R23E10> AstA-R1 RNAi v3400
AstA-R1 RNAi HMJ21471/+
UAS-Dcr2, R23E10> AstA-R1 RNAi HMJ21471
AstA neurons activation
f
25°C
31°C
Total Sleep (min)
UAS-dTrpa1/+
AstA-GAL4/+
AstA-GAL4>dTrpa1
LexAop-dTrpa1/+
65D05-LexA/+
65D05-LexA>LexAopdTrpa1

### Figure s4

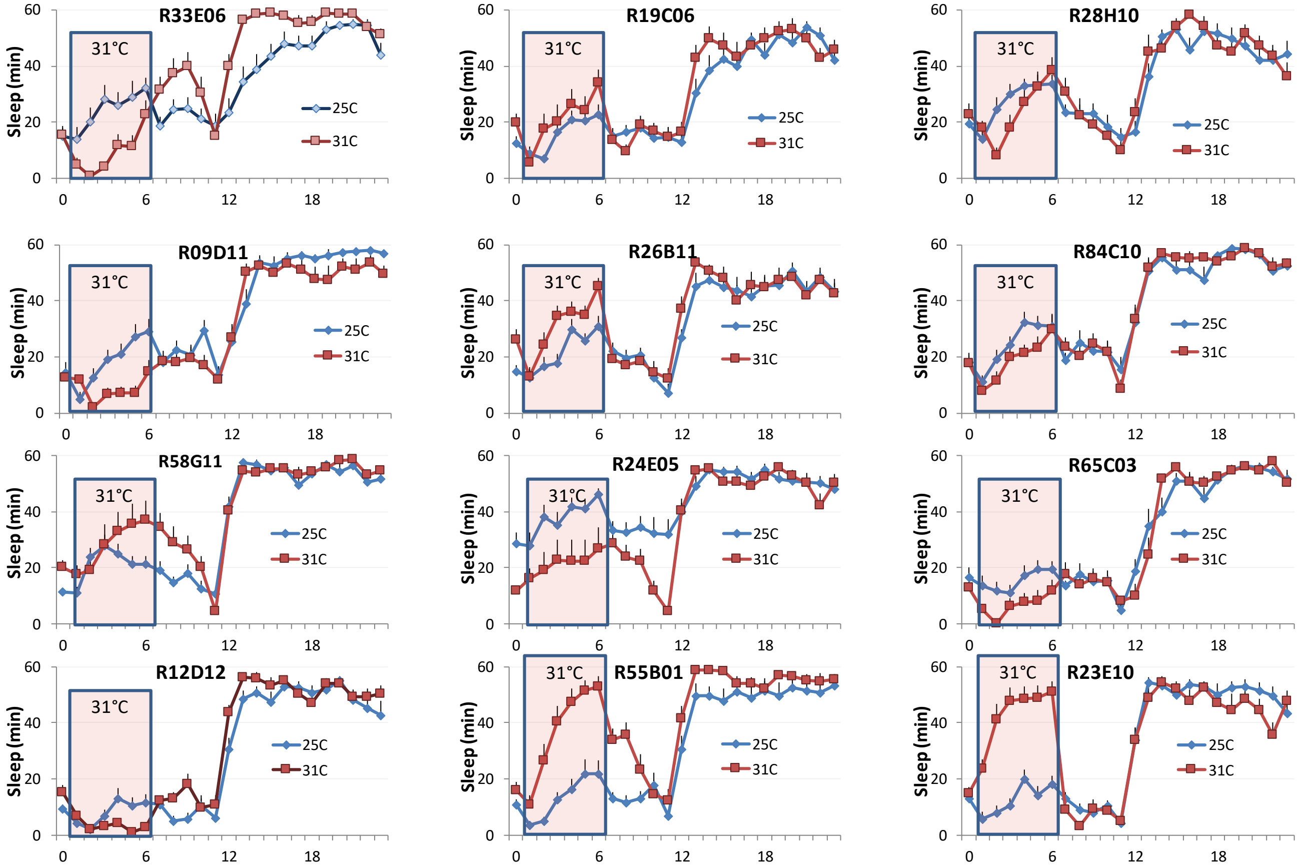
R33E06
R19C06
R28H10
R09D11
R26B11
R84C10
R58G11
R24E05
R65C03
R12D12
R55B01
R23E10
31°C
Sleep (min)
25C
31C
0
6
12
18
20
40
60

### Figure s5

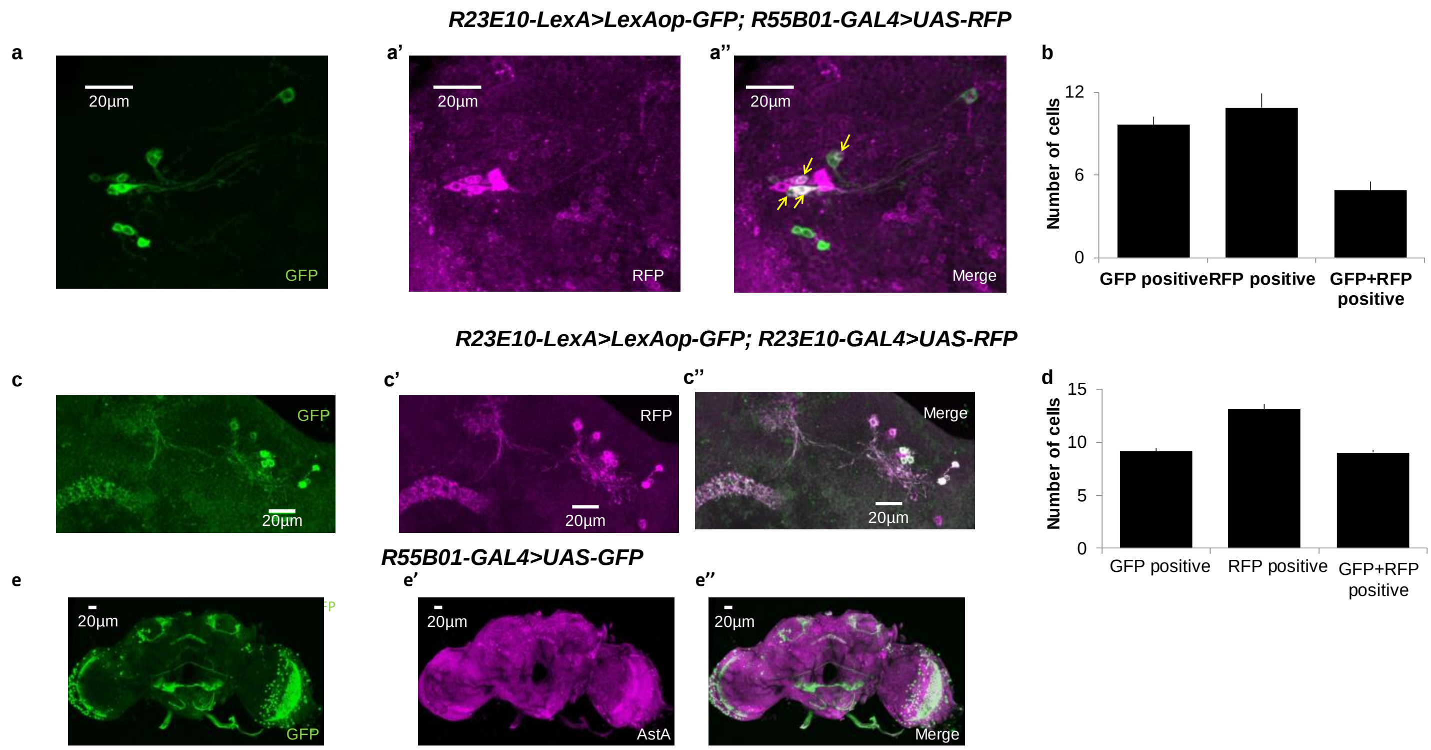
R23E10-LexA>LexAop-GFP; R55B01-GAL4>UAS-RFP
a
a’
a”
b
20µm
GFP
RFP
Merge
Number of cells
12
6
0
GFP positive
RFP positive
GFP+RFP positive
R23E10-LexA>LexAop-GFP; R23E10-GAL4>UAS-RFP
c
c’
c”
d
15
10
5
R55B01-GAL4>UAS-GFP
e
e’
e”
AstA

Figure s6

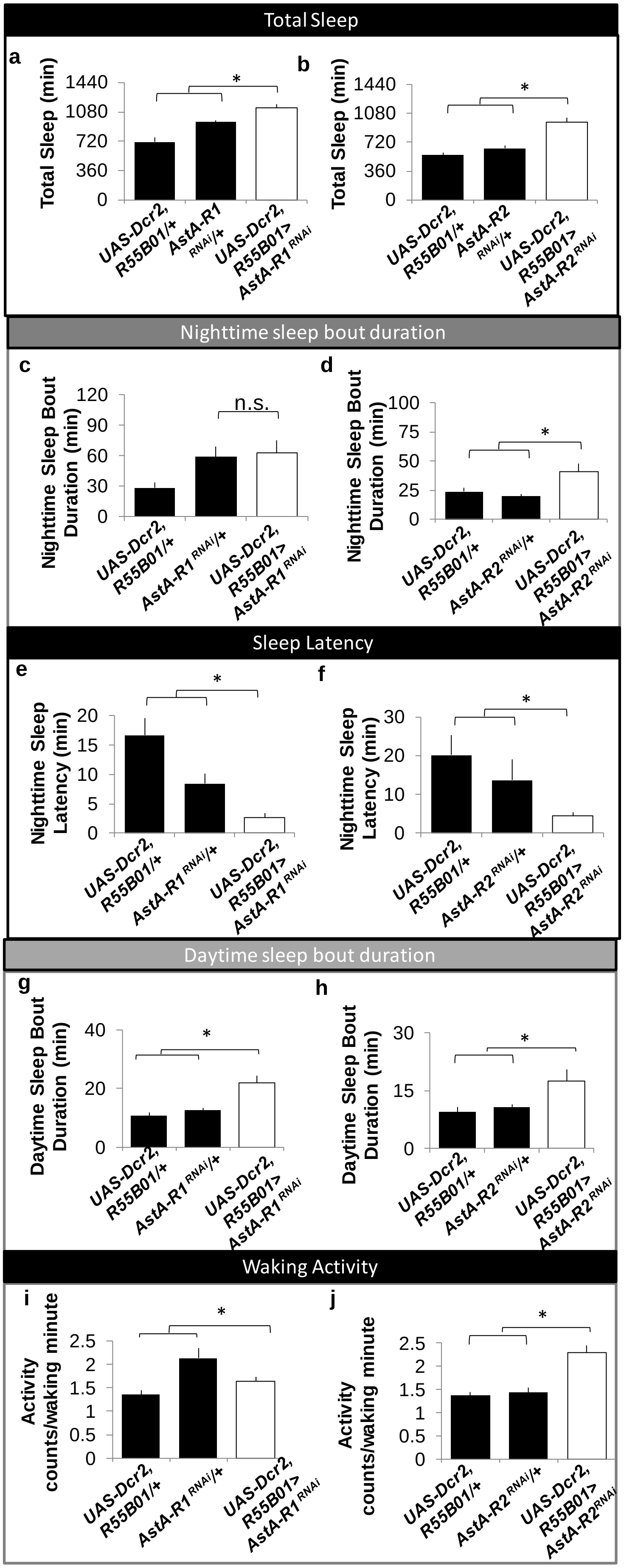

Total Sleep
a
Total Sleep (min)
1440
1080
720
360
0
*
UAS-Dcr2, R55B01/+
AstA-R1 RNAi/+
UAS-Dcr2, R55B01> AstA-R1 RNAi
b
Total Sleep (min)
*
UAS-Dcr2, R55B01/+
AstA-R2 RNAi/+
UAS-Dcr2, R55B01> AstA-R2 RNAi
Nighttime sleep bout duration
c
Nighttime Sleep Bout Duration (min)
120
90
60
30
0
n.s.
d
Nighttime Sleep Bout Duration (min)
100
75
50
25
0
*
Sleep Latency
e
Nighttime Sleep Latency (min)
20
15
10
5
0
*
f
Nighttime Sleep Latency (min)
30
20
10
0
*
Daytime sleep bout duration
g
Daytime Sleep Bout Duration (min)
40
20
0
*
h
Daytime Sleep Bout Duration (min)
30
15
0
*
Waking Activity
i
Activity counts/waking minute
2.5
2
1.5
1
0.5
0
*
j
Activity counts/waking minute
*

Figure s7

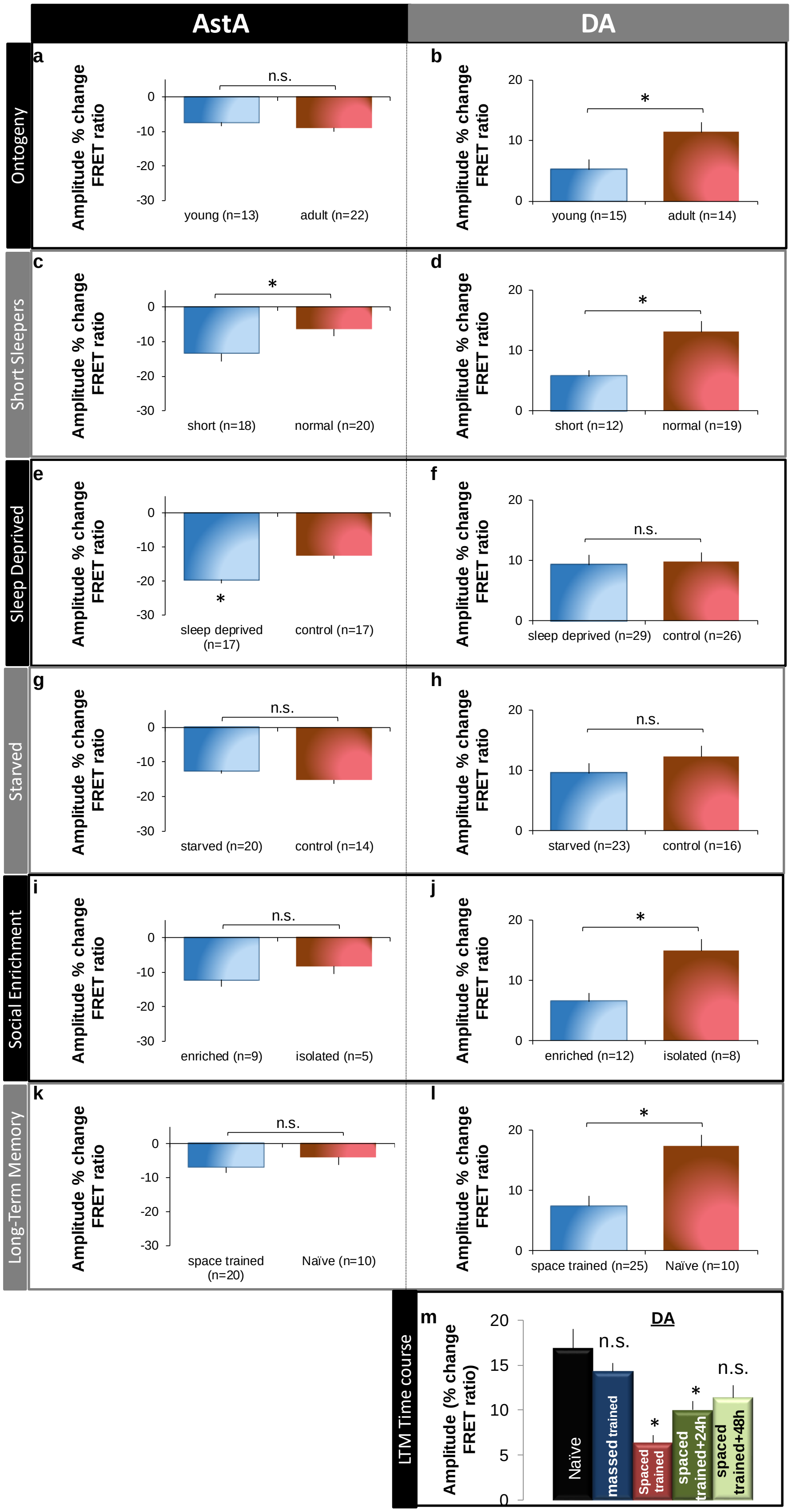
AstA
DA
Ontogeny
a
Amplitude % change FRET ratio
n.s.
young (n=13)
adult (n=22)
b
*
young (n=15)
adult (n=14)
Short Sleepers
c
*
short (n=18)
normal (n=20)
d
*
short (n=12)
normal (n=19)
Sleep Deprived
e
*
sleep deprived (n=17)
control (n=17)
f
n.s.
sleep deprived (n=29)
control (n=26)
Starved
g
n.s.
starved (n=20)
control (n=14)
h
n.s.
starved (n=23)
control (n=16)
Social Enrichment
i
n.s.
enriched (n=9)
isolated (n=5)
j
*
enriched (n=12)
isolated (n=8)
Long-Term Memory
k
n.s.
space trained (n=20)
Naïve (n=10)
l
*
space trained (n=25)
Naïve (n=10)
LTM Time course
m
Amplitude (% change FRET ratio)
DA
n.s.
n.s.
Naïve
massed trained
Spaced trained
spaced trained+24h
spaced trained+48h

Figure S8

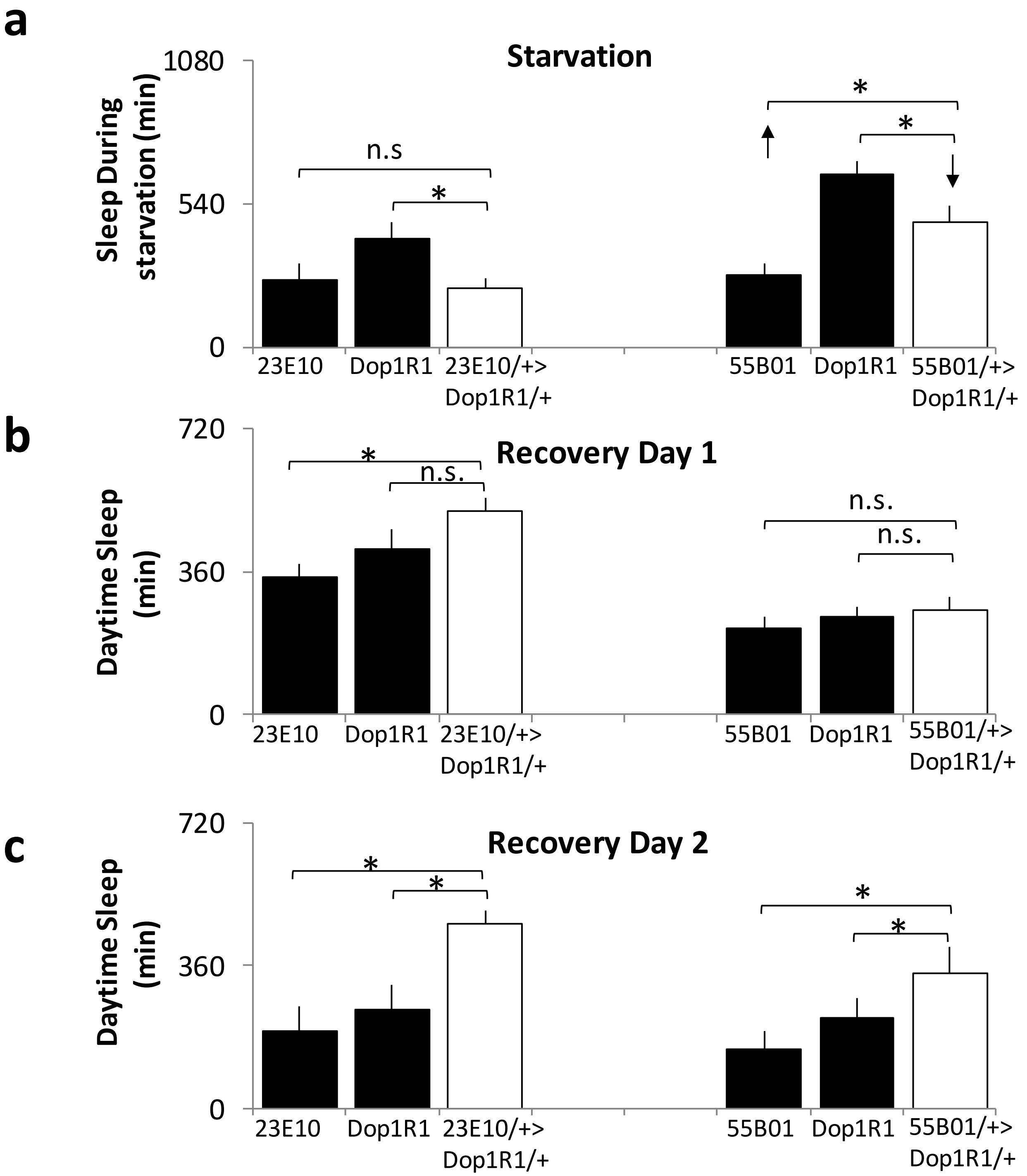

a
Starvation
Sleep During starvation (min)
1080
540
0
n.s
*
*
*
23E10
Dop1R1
23E10/+> Dop1R1/+
55B01
Dop1R1
55B01/+> Dop1R1/+
b
Recovery Day 1
Daytime Sleep (min)
720
360
0
*
n.s.
n.s.
n.s.
23E10
Dop1R1
23E10/+> Dop1R1/+
55B01
Dop1R1
55B01/+> Dop1R1/+
c
Recovery Day 2
Daytime Sleep (min)
720
360
0
*
*
*
*
23E10
Dop1R1
23E10/+> Dop1R1/+
55B01
Dop1R1
55B01/+> Dop1R1/+
